# The common analgesic paracetamol enhances the anti-tumour activity of decitabine through exacerbation of oxidative stress

**DOI:** 10.1101/2020.03.31.017947

**Authors:** Hannah J. Gleneadie, Amy H. Baker, Nikolaos Batis, Jennifer Bryant, Yao Jiang, Samuel J.H. Clokie, Hisham Mehanna, Paloma Garcia, Deena M.A. Gendoo, Sally Roberts, Alfredo A. Molinolo, J. Silvio Gutkind, Ben A. Scheven, Paul R. Cooper, Farhat L. Khanim, Malgorzata Wiench

## Abstract

The DNA demethylating agent 5-aza-2’-deoxycytidine (DAC, decitabine) has anti-cancer therapeutic potential, but its clinical efficacy is hindered by DNA damage-related side effects. Here we describe how paracetamol augments the effects of DAC on cancer cell proliferation and differentiation, without enhancing DNA damage. Firstly, DAC specifically upregulates cyclooxygenase-2-prostaglandin E_2_ pathway, inadvertently increasing cancer cell survival, while the addition of paracetamol offsets this effect. Secondly, combined treatment leads to glutathione depletion and ROS accumulation with oxidative stress further enhanced by DAC suppressing anti-oxidant and thioredoxin responses. The benefits of combined treatment are demonstrated here in head and neck squamous cell carcinoma (HNSCC) and acute myeloid leukaemia cell lines, further corroborated in a HNSCC xenograft mouse model and through mining of publicly available DAC and paracetamol responses. In summary, the addition of paracetamol could allow for DAC dose reduction, widening its clinical usability and providing a strong rationale for consideration in cancer therapy.

## Introduction

Aberrant DNA methylation patterns are common in most cancers, arise early in tumour development and are potentially reversible by hypomethylating agents [1]. The DNA demethylating agent 5-aza-2’-deoxycytidine (Decitabine or DAC) is a nucleoside analogue that incorporates into replicating DNA in place of cytosine where it traps and promotes the degradation of DNA methyltransferases (DNMTs) [2]. This results in two anti-cancer activities: methyl marks cannot be copied during DNA replication causing widespread DNA demethylation and adducts are formed in the DNA leading to DNA damage and apoptosis [2]. DNA demethylating drugs are thought to de-repress epigenetically silenced tumour suppressor genes as well as demethylate endogenous retroviruses (ERVs), triggering an antiviral immune response and cancer cell death [2-4]. DAC has been approved by the European Medicines Agency (EMA) for treatment of acute myeloid leukaemia (AML) [5, 6] while pre-clinical studies suggest it might also be effective in solid tumours [7]. However, the outcomes of clinical trials are highly variable, which is likely attributed to small sample sizes, lack of patient stratification, and inappropriate dosing and schedual [8].

Head and neck squamous cell carcinoma (HNSCC), the 6^th^ most common cancer worldwide, has 5-year survival rates of ≤40%, highlighting a pressing need for new therapies [9].The tumours originate from stratified squamous epithelium of the oral cavity and pharynx where the cells in the basal cell layer proliferate and replenish the suprabasal layers which undergo terminal differentiation [10]. Although DNA methylation aberrations are common in HNSCC [11] the clinical evaluation of DAC potential in HNSCC is very limited [12].

In solid tumours, DAC alone may not be curative, but favourable effects were observed for epigenetic agents combined with other chemo- and immune-therapies [8]. So far it has not been explored whether the response to DAC could be enhanced by other compounds, not traditionally used in cancer treatment. In the current study, a custom-built library of 100-commonly used, cost-effective, off-patent drugs [13] was investigated for their ability to sensitise HNSCC cells to DAC treatment. Of the drugs tested, paracetamol was identified to work in synergy with DAC.

Paracetamol (acetaminophen) is the most commonly used analgesic and antipyretic in both Europe and the United States, present on the market since the 1950s [14]. Paracetamol affects the cyclooxygenase (COX) pathway wherein arachidonic acid (AA) is metabolized to prostaglandin H_2_ (PGH_2_) by either constitutively expressed COX-1 (PTGS1) or inducible COX-2 (PTGS2) [15]. PGH_2_ is then rapidly converted, by respective prostaglandin synthases, into effector prostanoids (prostaglandins PGE_2_, PGF_2_; PGI_2_ and PGD_2_ or thromboxane TXA) and these work, in part, through metabolite-specific G-protein coupled receptors to activate downstream pathways [15]. Additionally, other AA-derived metabolites are produced either through the lipoxygenase (LOX) or the monooxygenase (cytochrome P450) pathways [16].

The COX-2-PGE_2_ axis is associated with inflammation, growth and survival and is thought to contribute to the ‘inflammogenesis of cancer’ [17]. Increased expression of COX-2 and production of PGE_2_ are found in many solid tumours, including HNSCC, and correlate with tumour stage, metastasis and worse clinical outcome whilst low levels are associated with better response to chemotherapy [17, 18]. This pathway has also been implicated in the development of chemoresistance and linked to repopulation of cancer stem cells [19]. Hence, COX-2 inhibitors have been tested for their therapeutic anti-cancer potential, showing protection against cancer development [18, 20]. However, the only research that suggests paracetamol may have therapeutic potential in established tumours used overdose concentrations of the drug, either alone or in combination with other chemotherapeutics [21-23].

Here we show that paracetamol can be used at clinically relevant concentrations to sensitize cancer cells (both HNSCC and AML) to DAC treatment, allowing for DAC dose-reduction.

## Results

### HNSCC cell lines show bimodal sensitivity to DAC treatment

To establish the potential of DAC as a therapeutic in HNSCC, relative cell viability was determined in four HNSCC cell lines and in normal human oral keratinocytes (HOK) after 96h of treatment (Fig 1A). HOK cells showed only moderate sensitivity to DAC treatment with no decrease in viability at clinically relevant concentrations and an IC_50_ of 8.93 μM. Interestingly, the four HNSCC cell lines could be divided into two distinct groups; DAC-sensitive (VU40T, IC_50_ of 2.17 μM and HN12, IC_50_ of 0.81 μM) and those with little (SCC040, IC_50_ of 10.61 μM) to no sensitivity (UDSCC2) (Fig 1A). This pattern was mirrored in the ability of DAC to demethylate DNA in the sensitive cell lines only (Fig 1B). These data suggest that the efficacy of DAC treatment is proportional to its ability to demethylate DNA, therefore likely dependent on cellular drug uptake, activation or retention.

**Figure 1.**
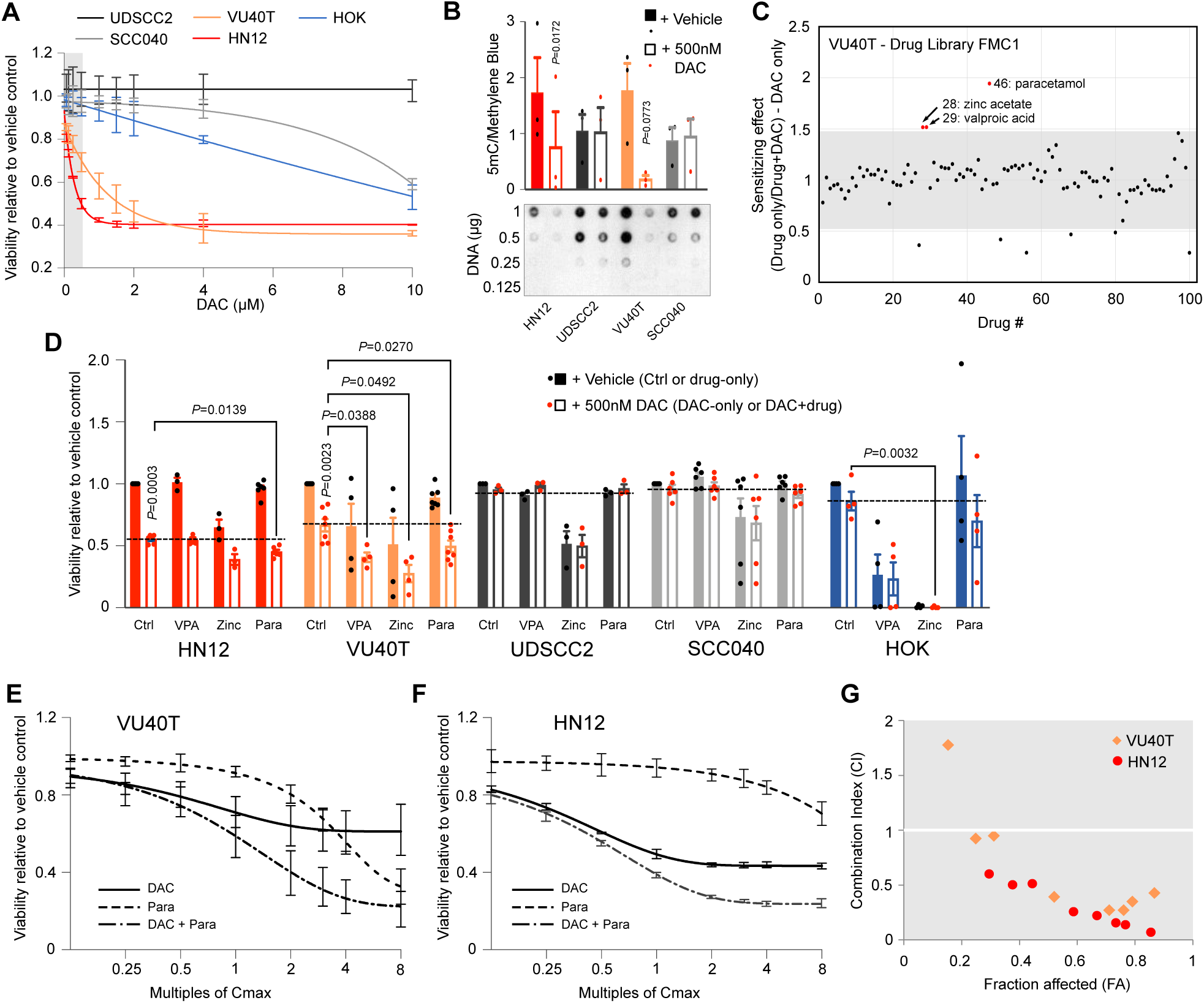
DAC therapeutic potential in HNSCC cell lines can be synergistically increased by co-treatment with paracetamol. **A**. Dose dependent cell viability in response to 96h DAC treatment in four HNSCC cell lines and normal oral keratinocytes (HOK). Grey box indicates clinically relevant doses. **B**.Global levels of DNA methylation (5mC) analysed by DNA dot blot in HNSCC cell lines +/- 500 nM DAC, 96h. The data were normalized to methylene blue staining and an example of 5mC dot blot is shown below. For each replicate the results for 1 and 0.5 μg of DNA were averaged. **C**. Scatterplot of the results from DAC sensitizing screen (full data in Appendix Fig S1) using 100 off patent drugs (Drug Library FMC1). Y axis represents increase in sensitivity, grey area shows ± 2SD. **D**. Cell viability in HNSCC cell lines and HOK cells treated for 96h with 132.3 μM paracetamol (Para), 601.7 μM valproic acid (VPA) and 323.4 μM zinc acetate (Zinc) +/- 500 nM DAC. The results are independent confirmation of data shown in Fig 1C and Appendix Fig S2. Dotted lines show the effect of 500 nM DAC alone. **E-G**. Synergy was determined using Chou-Talalay method. Cell viability data for VU40T (**E**) and HN12 (**F**) cells treated with fixed Cmax titrations of DAC (500 nM), paracetamol (132 μM) or DAC + paracetamol were used to calculate combination index (CI) (**G**) and dose reduction index (DRI) (Appendix Tables S1 and S2). CI <1 denotes synergy. DAC-sensitive cell lines are shown in reds, resistant – in greys, HOKs – in blue. Data information: In A-C and E-G n=3, in D n=7. In B statistical analysis was performed by paired two-tailed *t* tests. In D, for each cell line a matched One-Way ANOVA with Dunnett’s multiple comparison testing was used to compare DAC to DAC+drug. A separate paired *t* test was applied to compare no treatment with DAC. Values are displayed as means ± SEM. Significant p values are shown. Source data are available for this figure.

### Efficacy of DAC treatment can be synergistically increased with paracetamol

In patients with AML, a 5-day regimen of 20mg/m^2^ DAC gave a maximum plasma concentration (Cmax) of 107 ng/ml, equivalent to 469 nM [6, 24], while all HNSCC cell lines have an IC_50_ value greater than 500 nM. Therefore, one sensitive (VU40T) and one resistant (SCC040) cell line were subjected to DAC sensitising screen to establish whether the efficacy of DAC could be increased (Fig 1C, Appendix Fig S1). The cells were treated with 500 nM DAC with or without one of a panel of 100 off-patent drugs (drug repurposing library FMC1[13]) and compared to the drug-only control. In the resistant SCC040 cells, none of the drugs were able to sensitize the cells to DAC (Appendix Fig S1A). However, in the DAC-sensitive VU40T cell line, paracetamol, valproic acid (VPA) and zinc acetate further decreased cell viability (Fig 1C, Appendix Fig S1B). The paracetamol effect was replicated in the remaining cell lines: the efficacy of DAC was increased by paracetamol and zinc acetate in DAC-sensitive HN12 but not in DAC-resistant UDSCC2 cells (Fig 1D). Importantly, paracetamol alone did not alter the viability of HOK cells which were otherwise highly sensitive to both VPA and zinc acetate (Fig 1D). Therefore, DAC-paracetamol combination was further tested for synergy in VU40T and HN12 cells using the Chou-Talalay method [25]. The cell viability was assessed in response to each drug separately and in combination across eight constant-ratio matched titration of Cmax (500 nM for DAC and 132 μM for paracetamol) (Fig 1E-F). This analysis showed combination index (CI) values less than 1 at all bar the lowest concentration, demonstrating synergy of supra-additive nature (Fig 1G). In VU40T cells, the dose reduction index (DRI) indicated that almost 5 times less of each drug could be used when applied in combination (Appendix Table S1), allowing DAC dose reduction from 2.26 μM to the clinically relevant 450 nM [6, 24]. A similar reduction was observed for HN12 cells (Appendix Table S2). Although the synergistic effects are much stronger at higher concentrations, the Cmax was used throughout the study due to its clinical relevance.

### Combined DAC-paracetamol treatment augments the effects of DAC on cell metabolism, proliferation and markers of basal epithelial cells

In DAC-responsive VU40T cells the combined DAC-paracetamol treatment decreased cell number compared to DAC alone (Fig 2A). It was next investigated whether the underlying mechanisms involved reduced proliferation, altered cell cycle progression or increased cell death, possibly due to enhanced DNA damage.

**Figure 2.**
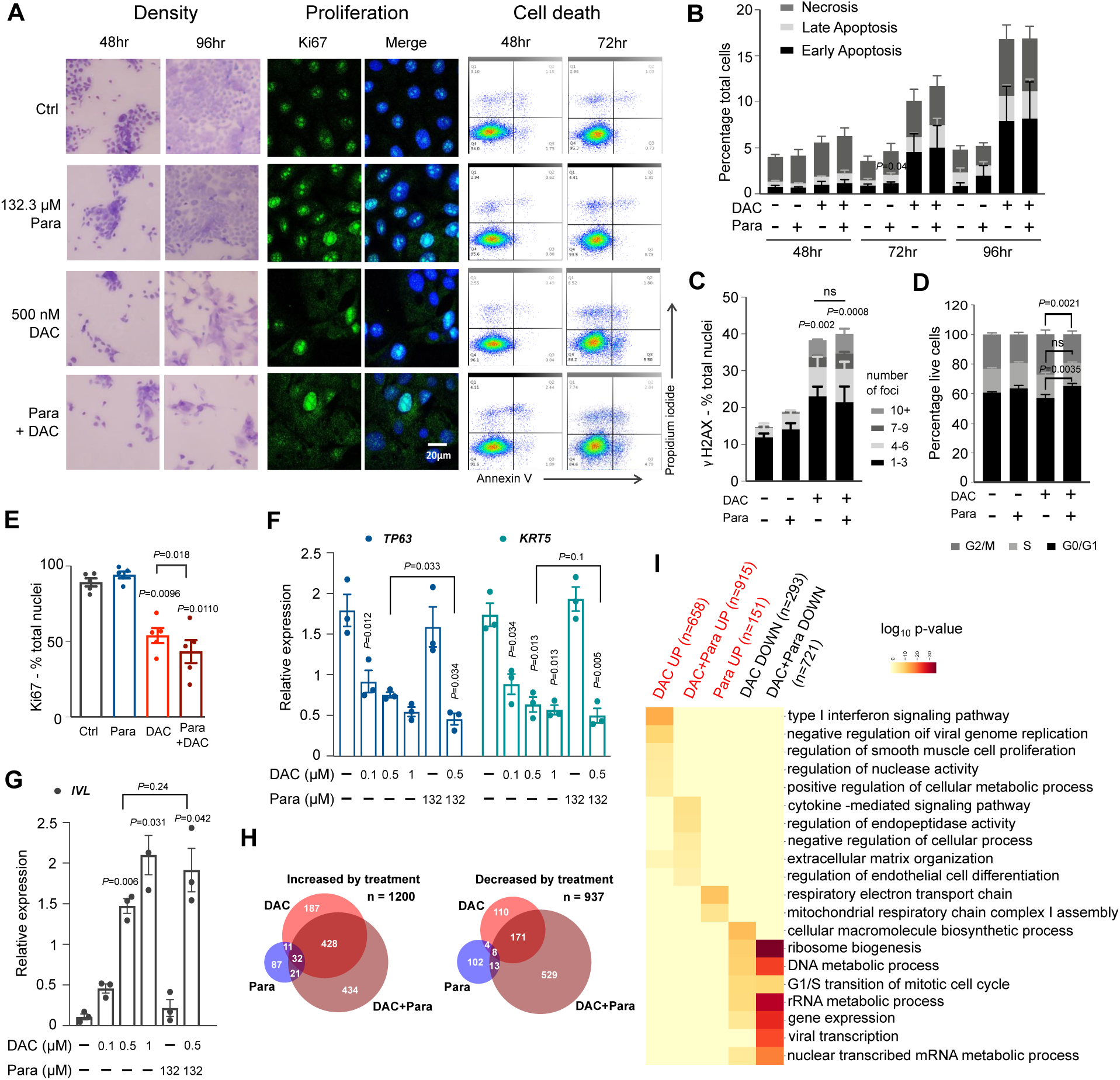
DAC-paracetamol combination enhances the effects of DAC on cell metabolism, proliferation and basal cell phenotype. **A**. Response of VU40T cells to paracetamol, DAC or both as indicated (representative images from 3 experiments). Left: Giemsa-Jenner staining (4x magnification); middle: Ki67 immunostaining; right: Annexin V and propidium iodine (PI) FACS analysis (healthy cells (bottom left); early apoptotic cells (bottom right); late apoptotic cells (top right) and necrotic cells (top left)). **B**. Proportion of VU40T cells undergoing cell death assessed by FACS following Annexin V and PI staining (examples shown in (A)). **C**. Percentage of VU40T nuclei with indicated numbers of γH2AX foci following indicated treatments. **D**. Cell cycle distribution in VU40T cells following indicated treatments. **E**. Proportion of VU40T nuclei positive for Ki67 immunostaining after indicated treatments (examples in (A)). **F**. qRT-PCR for *TP63* and *KRT5* normalized to *ACTB* in VU40T cells treated as indicated. **G**. qRT-PCR for *IVL* normalized to *ACTB* in VU40T cells treated as indicated. **H**.Venn diagrams showing overlap for the most upregulated (left, log_2_ fold change ≥1) and down-regulated (right, log_2_ fold change ≤-1) genes as compared to vehicle control. **I**. REVIGO-consolidated GO-BP pathways enriched for each indicated treatment. Top 5 terms from each group are included. Heatmap displays category scores as log_10_ p-value. Data information: Unless stated otherwise all treatments were for 96h with 500 nM DAC and 132.2 μM paracetamol. In B-D and F-G n=3, in E n=6. In B and D: Two-Way ANOVA with Dunnett’s correction was preformed to compare Ctrl with treatments and a separate Two-Way ANOVA with Sidak’s was performed to compare DAC with combined treatment. In C: matched Two-Way ANOVA with Tukey’s multiple comparison test was used to compare the distribution of foci number between each treatment group. In E-G: matched One-Way ANOVA with Dunnett’s multiple comparison test was used to compare all treatments to Ctrl. A separate paired two-tailed *t* test was used to compare DAC to DAC+paracetamol. Values are displayed as means ± SEM.

DAC is known to cause cell death and DNA damage [2]. As expected, 500 nM DAC increased both apoptosis and necrosis as shown by annexin V and propidium iodide staining but this was not altered by the addition of paracetamol (Fig 2A-B). Similarly, in VU40T cells, DAC treatment significantly increased DNA damage and double strand breaks as highlighted by an increase in the number of nuclei with γH2AX foci when compared to control; however, this too was not additionally enhanced by combined treatment (Fig 2C).

When compared to untreated cells none of the treatments resulted in change in cell cycle progression, however, the addition of paracetamol to DAC treatment led to fewer cells progressing to G2/M phase than for DAC alone (Fig 2D). This additional effect of paracetamol was also observed for cell proliferation. Treatment with 500 nM DAC reduced the number of Ki67 positive nuclei and combined treatment further decreased their number, whilst paracetamol alone had no effect (Fig 2A and E). Furthermore, markers for the basal cell layer (the only cell layer in stratified epithelium that normally contains dividing cells), *TP63* and keratin 5 (*KRT5*), were down-regulated by DAC in a dose-dependent manner and this was enhanced by DAC-paracetamol combined treatment (Fig 2F). Notably, TP63 is a key transcription factor in stratified epithelium [26] and its overexpression and/or amplification is observed in ∼ 30% of tumours in the TCGA HNSCC cohort (Appendix Fig S2). Conversely, involucrin (*IVL*), a marker for differentiated, suprabasal layers was upregulated upon DAC and even more so by DAC-paracetamol treatments (Fig 2G).

RNA sequencing performed on the DAC-sensitive VU40T cells demonstrated that DAC treatment substantially altered the transcriptome while paracetamol alone only resulted in modest changes (Fig 2H). However, addition of paracetamol to DAC treatment resulted in significantly greater changes in gene expression compared to DAC alone (Fig 2H). Six gene sets, up- and down-regulated by each treatment (Fig 2H, Appendix Table S3), were subjected to gene set enrichment analysis using GO Biological Processes [27] and consolidated in REVIGO [28] (Fig 2I, Appendix Table S4). Paracetamol treatment alone resulted in no significant enrichment for down-regulated gene sets; treatment primarily resulted in up-regulation of genes involved in respiratory electron transport chain. The profile upregulated by DAC was dominated by immune terms, especially interferon type I response. This was also evident in the combined treatment, although to a lesser extent. Interestingly, combined treatment was enriched for terms related to tissue development and differentiation (including ‘pharyngeal system development’ Appendix Table S4), confirming the possible change from basal cell-like to more differentiated epithelial cell phenotype. Genes down-regulated by both DAC and DAC+paracetamol showed similar ontology groupings related to DNA, protein and RNA metabolism but the enrichment was much stronger for the combined treatment (Fig 2I) due to higher number of differentially regulated genes (Fig 2H). This suggests that the addition of paracetamol to DAC significantly amplifies decreases in metabolism and DNA replication caused by DAC alone.

Therefore, although DAC alone has profound effects on proliferation, differentiation, cell death and DNA damage, the further reduction in viability observed after the addition of paracetamol appears to be associated with decreased proliferation and divergence from the basal cell-like phenotype.

### DAC treatment enhances the cyclooxygenase pathway, which is offset by paracetamol

Paracetamol is understood to act on the cyclooxygenase pathway, primarily through inhibition of COX-2 (PTGS2) [15]. Notably, a geneset for ‘Prostanoid biosynthetic process’ was enriched after both DAC and combined treatments (Appendix Table S4). Therefore, the effect of DAC on this pathway was further examined. In DAC-sensitive cell lines only, *PTGS2* RNA and protein levels were upregulated following DAC treatment and this was maintained after combined treatment (Fig 3A and B). A corresponding increase in the downstream product, prostaglandin E_2_ (PGE_2_), was observed in the DAC-sensitive VU40T cells but not in the resistant SCC040 cells (Fig 3C). PGE_2_ returned to basal levels by the addition of 132 μM paracetamol (Fig 3C) which indicates that paracetamol dampens DAC-induced COX-2 pathway activation. In this model DAC affects gene expression levels while paracetamol blocks protein function. In addition, expression of the PGE_2_ receptors *PTGER1* and *PTGER2* increased in the DAC-sensitive cells VU40T and HN12, respectively, whilst no significant changes in *PTGER1-4* expression occurred in the DAC-resistant cells UDSCC2 and SCC040 (Fig 3D and Appendix Fig S3A). Transcriptional activation of the COX-2-PGE_2_ pathway by DAC was evident in the RNA-seq data, together with COX-2-TXA_2_, whilst the PGF_2_, PGI_2_ and PGD_2_, pathways were not upregulated (Appendix Fig S3B and C). In summary, DAC treatment specifically upregulates many aspects of the COX-2-PGE_2_ pathway, inadvertently providing the cancer cells with growth and survival potential, while the addition of paracetamol offsets this effect (Fig 3E).

**Figure 3.**
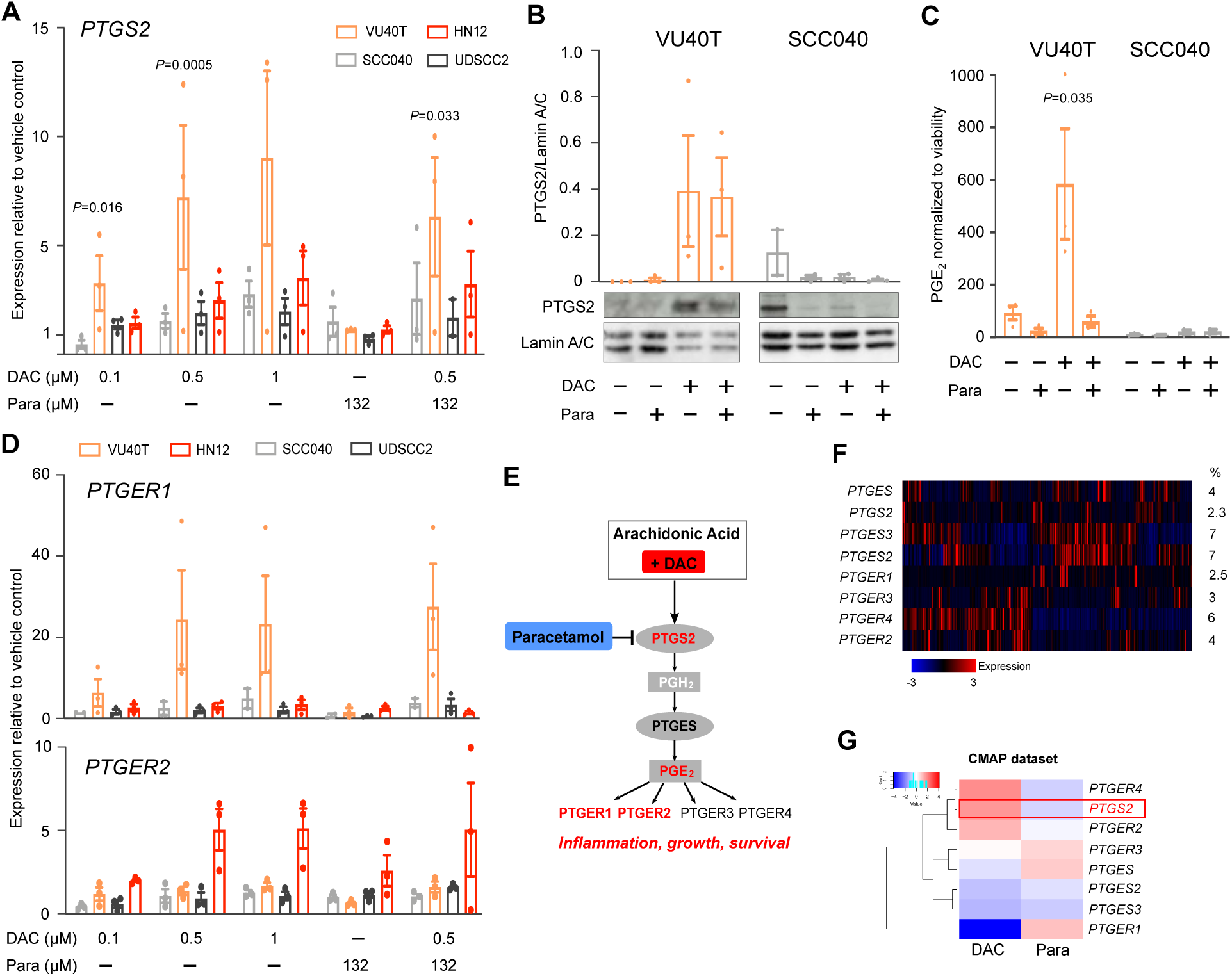
DAC treatment specifically activates COX-2-PGE_2_ pathway. **A**.qRT-PCR for *PTGS2* in HNSCC cell lines treated as indicated. Data are shown as relative to vehicle control (Ctrl=1). **B**.PTGS2 protein levels in VU40T and SCC040 cells. Graph represents the data from three experiments normalized to Lamin A/C. **C**.PGE_2_ concentration assessed by ELISA in media of VU40T and SCC040 cells treated for 96h as indicated. The data are normalized to cell viability. **D** qRT-PCR for PGE_2_ receptors, *PTGER1* and *PTGER2*, shown as in (A). **E**. Schematic of COX-2-PGE_2_ pathway with confirmed DAC effects shown in red. COX-2 catalyzes the conversion of arachidonic acid into prostaglandin H_2_. This is then converted into prostaglandin E_2_ which exerts its effects through G-protein coupled receptors. The full COX-2 pathway is shown in Appendix Fig S3B. **F**. COX-2-PGE_2_-related gene expression across 522 HNSCC tumour samples. Heatmap created in cBioPortal based on provisional HNSCC cohort (TCGA). % indicates fraction of tumours with alterations. **G**. Drug Perturbation Signature for genes of the COX-2-PGE_2_ pathway for Decitabine (DAC) and paracetamol (Para) using cMap dataset. Data information: Unless stated otherwise all treatments were for 96h with 500 nM DAC and 132.2 μM paracetamol and performed in three biological replicates. In A, B and D: for each cell line a matched One-Way ANOVA with Dunnett’s multiple comparison test was used to compare all treatments to Ctrl; a separate paired two-tailed *t* test was used to compare DAC to DAC+paracetamol. In C: for each cell line an ordinary One-Way ANOVA with Sidak’s correction was applied to compare treated groups with Ctrl. Values are displayed as means ± SEM. Only significant p-values are shown. Source data are available for this figure.

There is a possibility that blocking the COX pathway with paracetamol could shunt AA towards the LOX pathway also leading to increased survival potential [29]. DAC treatment altered gene expression of enzymes involved in the LOX pathway by both up-(*ALOX5* and *ALOX15B*) and down-(*LT4H* and *ALOX12*) regulating them (Appendix Fig S3D). However, the secretion of cysteinyl leukotrienes and leukotriene B_4_ remained below levels detectable by ELISA in all experimental conditions in VU40T and SCC040 cells. Therefore, there is no indication for LOX pathway compensation following COX-2 inhibition by paracetamol in the cell lines examined, although the involvement of other metabolites cannot be excluded (Appendix Fig S3E).

### DAC treatment complements cancer-related activation of COX-2-PGE_**2**_ **pathway**

Activation of the COX-2-PGE_2_ pathway has been previously indicated in both HNSCC and other cancer types [17, 18]. In the most recent TCGA cohort, 29% of HNSCC tumours (150/520 patients) have at least one component of COX-2-PGE_2_ pathway transcriptionally activated: mostly through upregulation of PGE_2_ synthases or PGE_2_ receptors (Fig 3F). Similar activation can be observed in other cancers (Appendix Table S5). However, over-expression of *PTGS2* itself is relatively rare (2.3% in HNSCC, Fig 3F), potentially serving as a limiting factor in the full activation of the pathway. Therefore, the DAC-induced upregulation of *PTGS2* could remove this limitation and counteract the anti-tumour effects of DAC. This mechanism could be applicable to other tumour types; *PTGS2* up-regulation by DAC was detected by Drug Perturbation Signatures (Fig 3G) from the cMAP dataset [30] indicating the wider potential of DAC-paracetamol co-treatment.

### Combined treatment mimics the effects of paracetamol overdose and depletes glutathione levels in HNSCC cells

If COX-2-PGE_2_ pathway alterations were to be solely responsible for the synergy between DAC and paracetamol, other COX inhibitors should have a similar effect. However, neither the generic COX inhibitor ibuprofen, nor the COX-2 specific inhibitor valdecoxib sensitized HNSCC cells to DAC treatment (Fig 4A, Appendix Fig S4), despite valdecoxib being able to reduce DAC-stimulated PGE_2_ production to a similar extent as paracetamol (Fig 4B). This suggests an additional, paracetamol-specific, mechanism accounts for the synergistic relationship between DAC and paracetamol.

**Figure 4.**
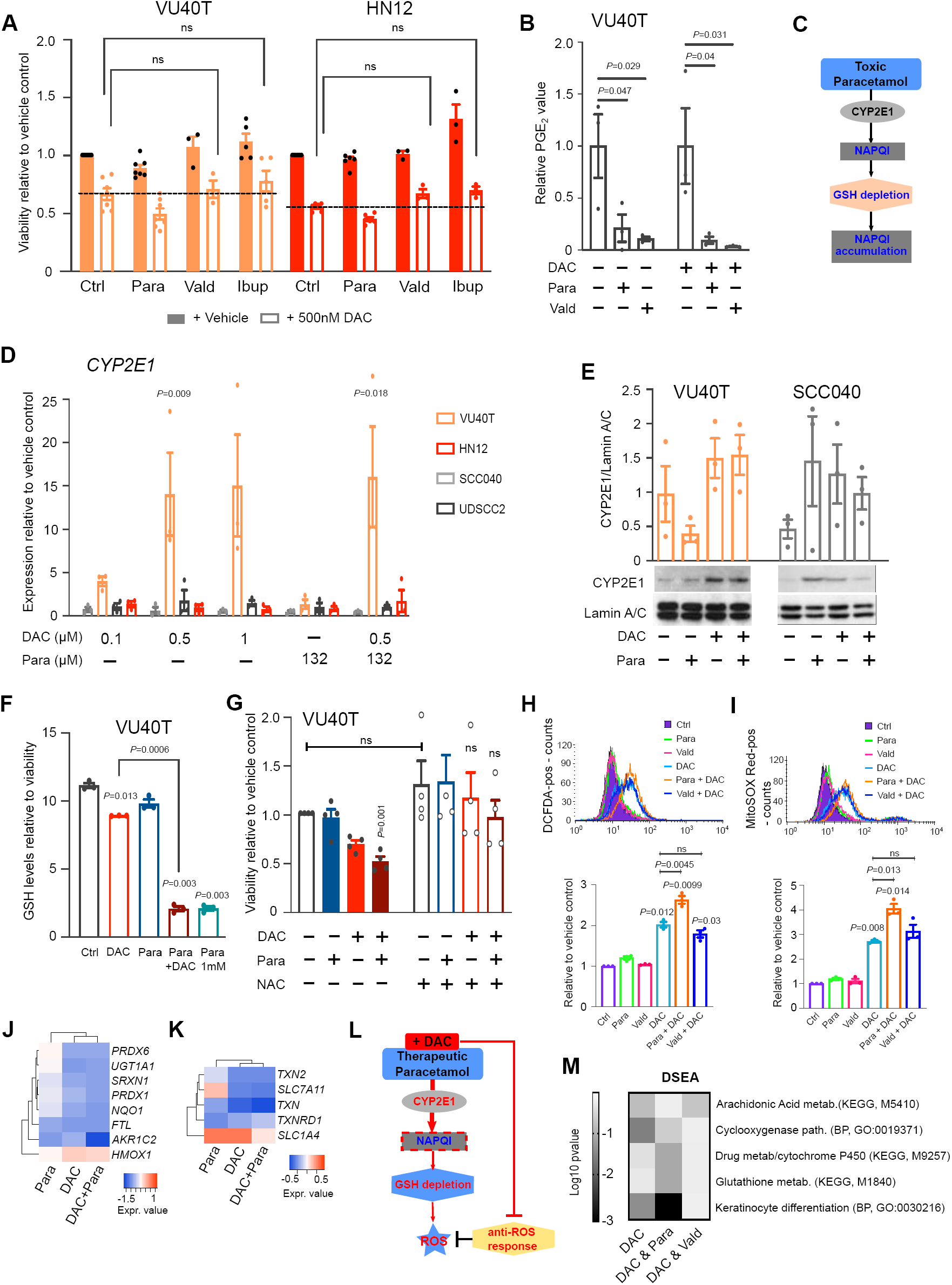
Combined DAC-paracetamol treatment mimics the effects of paracetamol overdose, depletes glutathione levels and leads to oxidative stress. **A**. Cell viability in VU40T and HN12 cells treated for 96h with 132.3 μM paracetamol, 10 μM valdecoxib or 193.9 μM ibuprofen +/- 500nM DAC. Dotted line shows the effect of DAC alone. **B** .PGE_2_ concentration in the media of VU40T cells treated with DAC with or without paracetamol or valdecoxib shown as relative to either vehicle control (left) or DAC only treatment control (right). **C**. Schematic of paracetamol overdose. A fraction of paracetamol is converted into the toxic metabolite NAPQI, and detoxified by GSH. When paracetamol is taken in excess GSH stores deplete and NAPQI accumulates. **D**. qRT-PCR for *CYP2E1* in HNSCC cell lines treated as indicated and shown relative to vehicle control (Ctrl=1). **E**. CYP2E1 protein levels in VU40T and SCC040 cells. The graph represents the data from three experiments normalized to Lamin A/C. **F**. Glutathione (GSH) levels in VU40T cells treated with DAC, paracetamol or both, normalized to cell viability. 1 mM paracetamol was used as a positive control. **G**. NAC rescue experiment. VU40T cells were pre-treated for 48h with 2.5 mM NAC followed by 96h of DAC, paracetamol or combined treatments and cell viability was assessed in comparison to control without NAC (left panel). **H**. Levels of intracellular ROS (DCFDA staining and FACS) in VU40T cells treated for 72h as indicated. Upper figure: representative FACS profile; lower graph: geometric mean normalized to vehicle control. **I**. Levels of mitochondrial superoxide were assessed as in (H) by MitoSOX Red staining. **J**. Gene expression (RNA-seq data) of direct responders to oxidative stress shown as heatmap of log_2_ fold change values after indicated treatments. **K**. Gene expression of cystin/TXN/TXNRD1 system components shown as in (J). **L**. Schematic of proposed mechanism underlying DAC-paracetamol synergy: mimicry of paracetamol overdose leading to GSH depletion and oxidative stress. The effects of DAC are shown in red and paracetamol contribution – in blue. **M**. Drug Set Enrichment Analysis for DAC, DAC + paracetamol, and DAC + valdecoxib combinations. Log_10_ p-values for selected pathways of interest (KEGG or GO BP genesets) are shown as heatmap. Data information: Unless stated otherwise all treatments were for 96h with 500 nM DAC and 132.2 μM paracetamol, n=3 for A-B, D-F, H-I and n=4 for G. In A: for each cell line a mixed-effect analysis with Dunnett’s correction was used to compare DAC to DAC+Vald and DAC+Ibup samples. In B, D-F: One-Way ANOVA with Dunnett’s correction was used to compare treatments with Ctrl. In D and F: a separate paired two-tailed *t* test was used to compare DAC to DAC+paracetamol. In G: for each group (+vehicle, +NAC) a matched One-Way ANOVA with Dunnett’s was used to compare treatments with Ctrls; additionally, a paired two-tailed *t* test was used to compare Ctrl with NAC. In H-I: a matched One-Way ANOVA with Dunnett’s correction was used to compare all groups to Ctrl; a separate ANOVA was used to compare DAC to DAC+Para and DAC+Vald. Values are displayed as means ± SEM. Only significant p values are shown. Source data are available for this figure.

Previous work on the anti-cancer therapeutic potential of paracetamol involved toxic doses of the drug [21-23]. Efficacy was attributed to an accumulation of toxic metabolite of paracetamol, N-acetyl-p-benzoquinone-imine (NAPQI), resulting in glutathione depletion [21-23] (Fig 4C). By comparison, our current work was performed using a safe, clinically achievable serum concentration of paracetamol (132 μM). However, in DAC-sensitive VU40T cells, DAC treatment caused an increase in the CYP450 enzyme, *CYP2E1*, thought to be involved in the conversion of paracetamol into NAPQI (Fig 4D-E). In addition, increased expression of the majority of CYP enzymes-encoding genes was observed in RNA-seq dataset and some of these could also enhance paracetamol-NAC conversion in the absence of the specific *CYP2E1* up-regulation (Appendix Fig S4B). Assessing NAPQI levels is unreliable due to its highly reactive nature [31]. However, combined treatment led to a dramatic reduction in GSH levels, a surrogate marker often used, and the effect was equivalent to a high dose (1mM) paracetamol (Fig 4F). Furthermore, N-acetyl cysteine (NAC) is clinically used as an antidote to treat paracetamol overdose by replenishing GSH and preventing NAPQI accumulation [21]. In VU40T cells, 48h pre-treatment with 2.5 mM NAC led to increased cell viability in all conditions, but significantly and to the greatest extent in the DAC+paracetamol treated cells, where viability was restored to the control level (Fig 4G). Therefore, the combined treatment mimics the effects of paracetamol overdose at therapeutic concentrations and depletes GSH stores in tumour cells.

GSH is an intracellular antioxidant that acts primarily as a reactive oxygen species (ROS) scavenger [32]. In agreement with this, DAC-paracetamol co-treatment significantly increased both intracellular ROS (Fig 4H) and mitochondrial superoxide (Fig 4I) as compared to DAC alone. This was specific for DAC co-treatment with paracetamol and not valdecoxib (Fig 4H-I). Accumulation of ROS typically triggers an anti-oxidant response which can protect cancer cells treated with chemotherapeutics [33]. However, upon both DAC and DAC-paracetamol treatment the majority of genes described as direct anti-oxidant responders [34] are down-regulated (Fig 4J). Therefore, the transcriptional anti-ROS response is impaired by DAC treatment which leads to further exacerbation of oxidative stress and decreased cancer cell survival. Finally, it has been shown that keratinocytes can tolerate GSH depletion as long as the cysteine pools (uptake of cystine by SLC7A11 or cysteine by SLC1A4) and the thioredoxin reductase system (TXN/TXNRD) are functional [35]. Again, DAC downregulated the majority of these genes which would inhibit this compensating mechanism (Fig 4K).

To summarize, in cancer cells DAC-paracetamol co-treatment mimics the mechanisms of paracetamol overdose, whilst cell adaptation to oxidative stress is impaired by DAC (Fig 4L). The specificity of the DAC-paracetamol interaction is further supported through Drug Set Enrichment Analysis (DSEA). The DSEA algorithm aims to identify the mechanisms of action shared by a set of drugs [36]. Indeed, DAC and paracetamol share significant enrichment for COX, cytochrome P450 and GSH metabolism pathways, while this was not observed for DAC and valdecoxib combination (Fig 4M, Appendix Fig S5). Interestingly, the analysis also indicated high enrichment for genes attributed to keratinocyte differentiation (Fig 4M).

### DAC-paracetamol affected pathways contribute to HNSCC patients’ survival

The mechanisms underlying the efficacy of DAC-paracetamol co-treatment identified so far are multifactorial (Fig 5A). Firstly, the COX-2-PGE_2_ pathway is specifically upregulated by DAC, potentially improving cancer cells survival. Indeed, when upregulated, this pathway negatively affects HNSCC patient survival (Fig 5B). This is specific to COX-2-PGE_2_ but not to other COX pathways or to the LOX pathway (Appendix Fig S6A and B). The addition of paracetamol to DAC treatment counteracts the induction of this pro-survival mechanism. Secondly, combined treatment leads to depletion of glutathione stores followed by oxidative stress, both of which restrict tumour growth and improve the efficacy of anti-cancer drugs [37]. Therefore, in the proposed scenario, GSH is required to maintain cancer cell fitness. In agreement with this, genes involved in maintaining GSH stores are linked to poorer patients’ survival in the TCGA cohort of HNSCC tumours (Fig 5C) as well as other cancers (Appendix Table S6). Thirdly, DAC treatment transcriptionally down-regulates anti-oxidant and thioredoxin responses known to prevent cellular adaptation to oxidative stress and protection from oxidative damage. Again, upregulation of these pathways, particularly the thioredoxin response, correlates with decreased survival of HNSCC patients (Fig 5D). These results suggest that the DAC-paracetamol combination targets pathways and genes of clinical importance in HNSCC patients.

**Figure 5.**
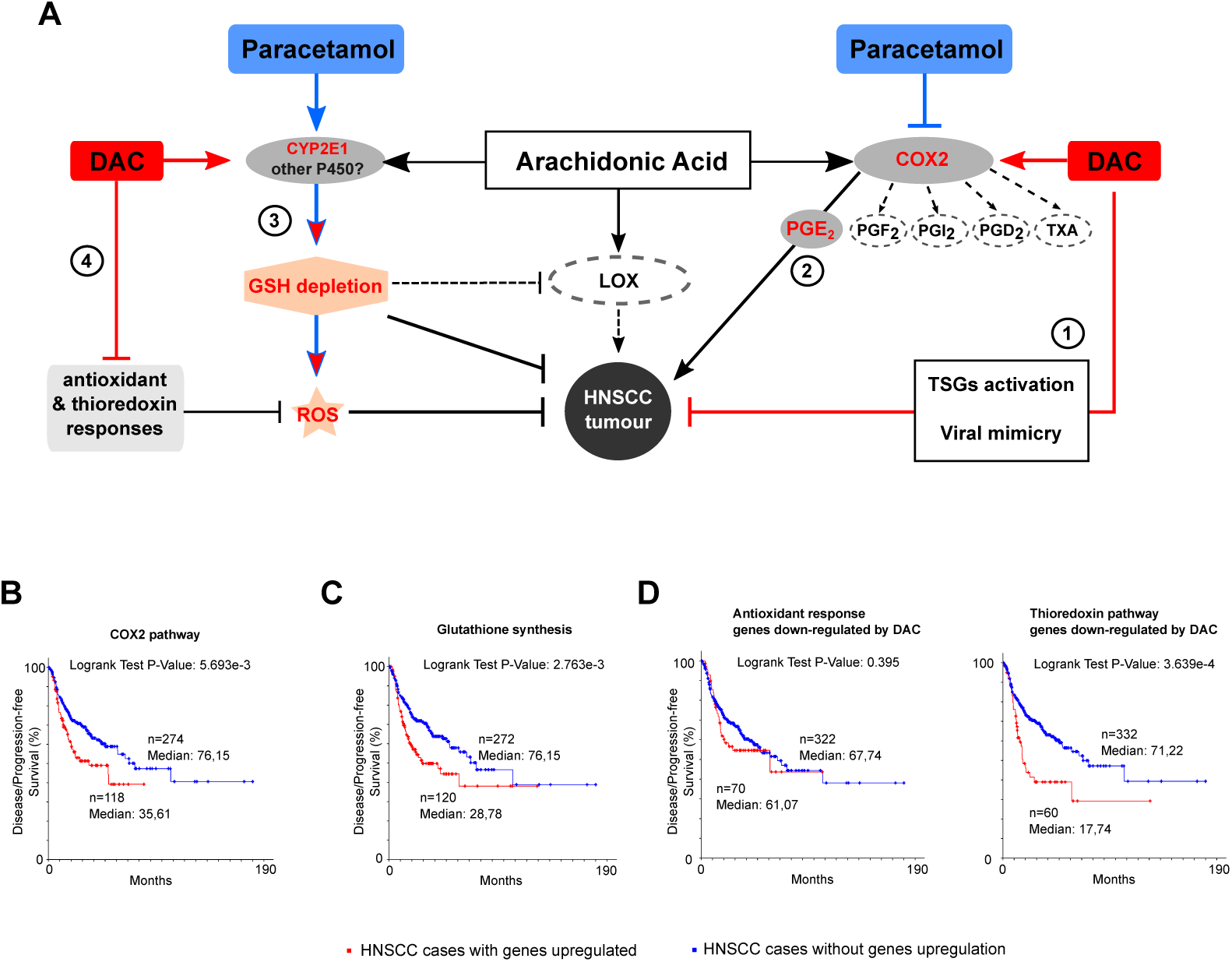
Combined effects of DAC and paracetamol on AA metabolism and HNSCC patients’ survival. **A**. Arachidonic acid (AA) is metabolized to eicosanoids through COX, LOX, and cytochrome P450 monooxygenase pathways. In addition to DAC limiting cancer cell growth through either activation of tumour suppressor genes (TSGs) or viral mimicry (1), it inadvertently activates COX-2 PGE_2_ pathway (2), which is contradicted by paracetamol. DAC also upregulates *CYP2E1* which, in the presence of paracetamol, leads to glutathione depletion and ROS accumulation, both enhanced by combined treatment (3). Simultaneously, DAC downregulates transcription of genes involved in antioxidant and thioredoxin responses, preventing cancer cells from developing adaptation to oxidative stress and protection from oxidative damage (4). GSH depletion has also potential to limit the production of LOX pathway metabolites dependent on GSH transferases. **B-D**. Survival curves for 392 HNSCC patients with data available in TCGA provisional cohort with or without upregulation of genes (z-score ≥2.0) involved in indicated pathway. For gene sets see Appendix Materials and Methods. Kaplan Meier Estimates were plotted using cBioPortal. (**B**) Disease/ Progression-free survival curves for genes involved in COX-2-PGE_2_ pathway. (**C**) Disease/Progression-free survival curves for genes involved in glutathione synthesis. (**D**) Disease/Progression-free survival curves for genes involved in antioxidant (left) and thioredoxin (right) responses.

### *In vivo* potential of DAC-paracetamol combination treatment

An NSG mouse xenograft model using human HNSCC FaDu cells previously used in drug efficacy studies [38] was utilized to assess the DAC-paracetamol treatment *in vivo*. Like HN12 and VU40T, FaDu cells showed high supra-additive effects for combined treatment (Fig 6A and B). Mice were injected with FaDu cells and the treatments (DAC, paracetamol, DAC+paracetamol or vehicle (PBS)) were administered 5 days a week (Fig 6C). After the first two weeks, tumours in the control and paracetamol-treated groups reached the maximum permissible size, while DAC alone and DAC+paracetamol groups were treated for another week (Fig 6C). Due to the strong initial response to DAC, the treatment was carried out with reduced DAC concentration (0.2 mg/kg). Although DAC alone showed a strong anti-tumour effect, the DAC-paracetamol treated tumours remained consistently smaller throughout the duration of the treatment (Fig 6C and D). No tumours from the DAC-paracetamol treated group exceeded 300 mm^3^ (Fig 6E) and 5/6 animals survived until the end of the experiment at which point they started losing weight (Appendix Fig S7). Transcript analysis of RNA extracted from tumour tissue showed alterations consistent with the mechanism of DAC-paracetamol synergy: upregulation of *PTGS2* and *CYP2E1* (Fig 6F) and down-regulation of *TP63* and *KRT5* (Fig 6G).

**Figure 6.**
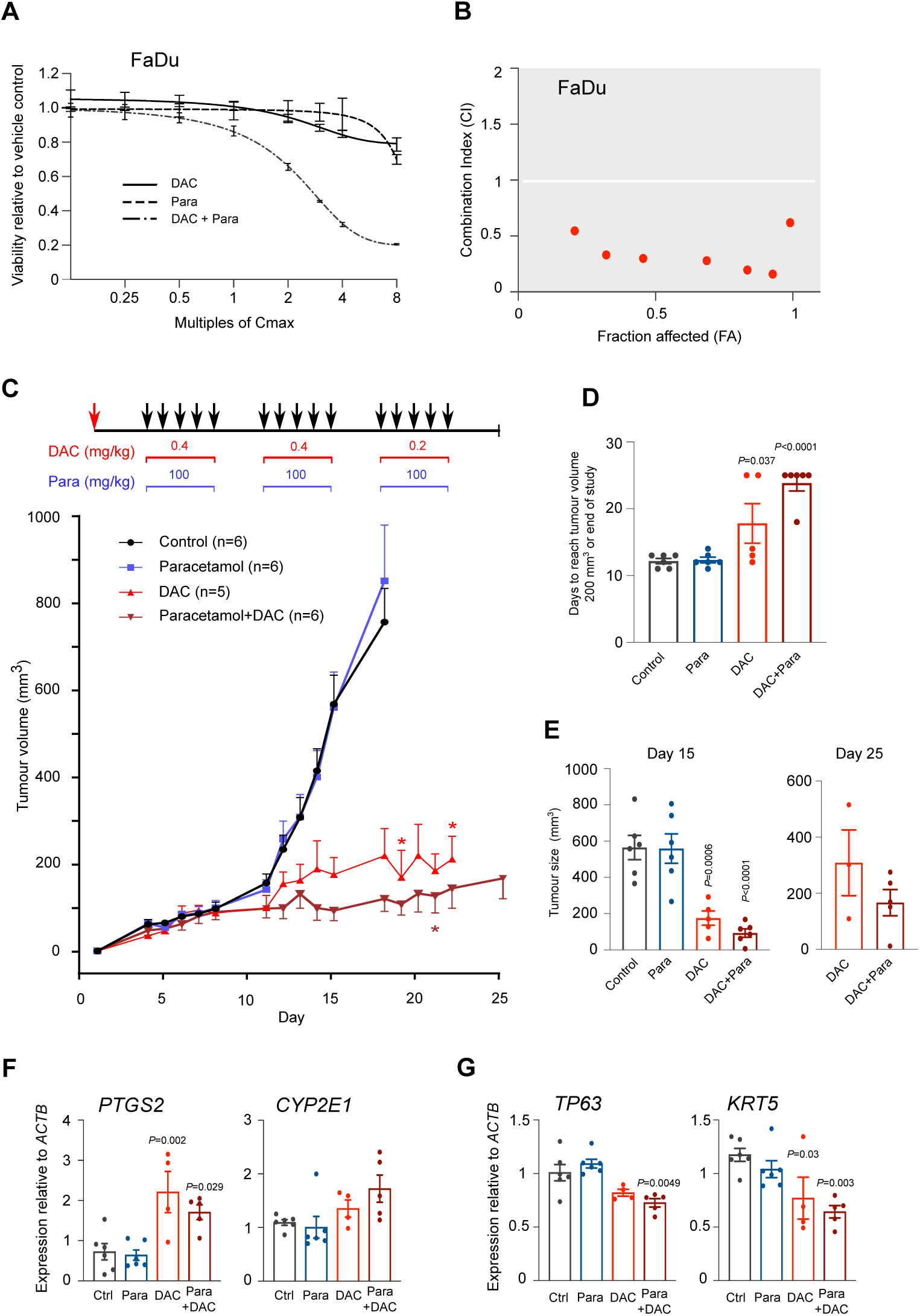
*In vivo* potential of DAC-paracetamol combination. **A-B**. DAC-paracetamol synergy assessed in FaDu cells (as in Fig 1E-G); cell viability after treatment with fixed Cmax titrations of drugs (**A**) resulting in combination index (CI) shown in (**B**). CI value less than 1 denotes synergy. **C**. Tumour growth of FaDu cells engrafted into NSG mice (Day 1, red arrow) treated as indicated (black arrows). **D**. Days to reach tumour size of 200 mm^3^. Tumours below 200 mm^3^ at the end of the experiment are counted as Day 25. **E**. Tumour size at Day 15 and at the end of the study. **F**. qRT-PCR for *PTGS2* and *CYP2E1* expression in tumour tissue. **G**. qRT-PCR for *TP63* and *KRT5* expression in tumour tissue. Data information: In A-B n=3. Mice groups: Ctrl (n=6), paracetamol (n=6), DAC (n=5), DAC+paracetamol (n=6). In D-G: a non-matched One-Way ANOVA with Dunnett’s correction was used to compare treatments with control. To compare DAC and DAC+Para an unpaired two-tailed *t* test was used. Values are displayed as means ± SEM. Only significant p values are shown.

### Synergistic effects of DAC-paracetamol co-treatment are present in AML cell lines

Currently, DAC only has EMA approval in the treatment of AML [6]. Therefore, combined treatment of DAC plus paracetamol was tested in two AML cell lines, SKM-1 and HL-60. The cells were treated for 72h and the drugs were found to work synergistically in both of them (Fig 7A-C). The combined effect was even more apparent after prolonged culture, which aimed to mimic the DAC treatment regimen in AML patients [24] where the cells were treated for 72h followed by 21 days withdrawal over four cycles (Fig 7D). The benefits of DAC-paracetamol treatment were more pronounced with each consecutive cycle and by the time the cells reached the fourth cycle they were still responding well to combined treatment while becoming resistant to DAC alone (Fig 7D).

**Figure. 7.**
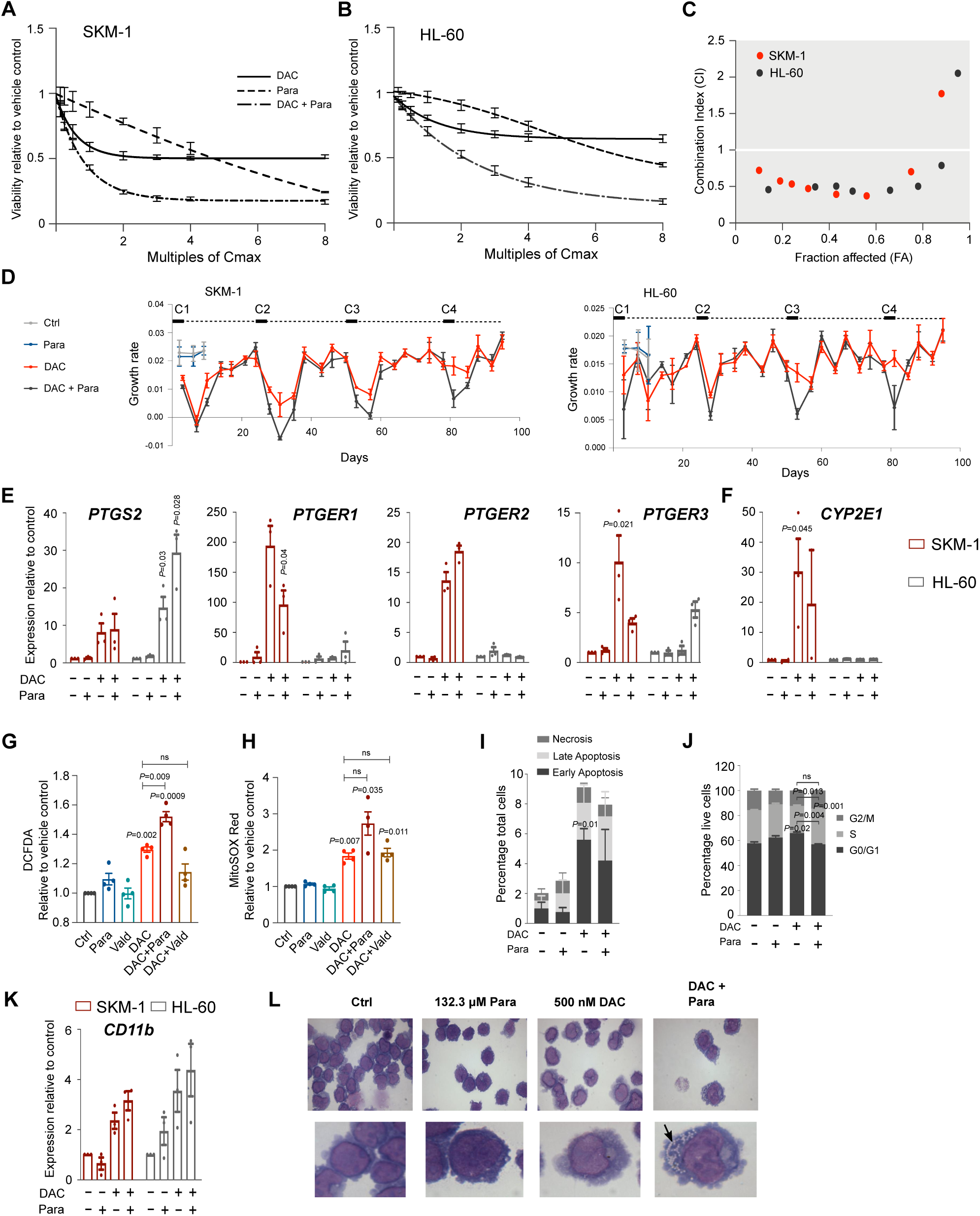
Potential of DAC-paracetamol combined treatment in AML. **A-C**. DAC-paracetamol synergy determined in AML cell lines. Cell viability of SKM-1 (**A**) and HL-60 (**B**) treated as indicated for 72h. Combination index (CI) shown in (**C**). **D**. Long term effects of DAC and DAC-paracetamol combined treatment on growth rates in SKM-1 (left) and HL-60 (right) cells: 4 cycles (C1-C4, 72h treatment as indicated followed by 21 day withdrawal). Vehicle and paracetamol controls are shown for the first 10 days. **E**. qRT-PCR for *PTGS2, PTGER1, PTGER2* and *PTGER3* genes in SKM-1 and HL-60 cells after indicated treatments and shown relative to control. **F**. qRT-PCR for *PTGS2* shown as in (E). **G**. Levels of intracellular ROS measured by DCFDA staining and FACS in SKM-1 cells treated with indicated drugs and their combinations. Vald, 10 μM valdecoxib. **H**. Levels of mitochondrial superoxide assessed by MitoSOX Red staining and FACS in SKM-1 cells as in (G). **I**. Proportion of SKM-1 cells undergoing cell death assessed by FACS following Annexin V and PI staining. **J**. Cell cycle distribution of SKM-1 cells after of DAC, paracetamol or combined treatment. **K**. qRT-PCR for *CD11b* gene in SKM-1 and HL-60 cells after indicated treatments. **L**. Giemsa-Jenner staining in SKM-1 cells. Upper: 100x magnification; lower: image zoomed to approximately one cell. Vacuole formation in cytoplasm indicated by black arrow. Data information: Unless stated otherwise all treatments were for 72h with 500 nM DAC and 132.2 μM paracetamol and n=3. In E-H and K: a matched One-Way ANOVA with Dunnett’s correction was applied to compare treatments with control; in G-H a separate ANOVA was used to compare DAC to DAC+Para and DAC+Vald. In I-J: for each group a matched Two-Way ANOVA with Dunnett’s was performed to compare treatments with control; a separate Two-Way ANOVA with Sidak’s was used to compare DAC with DAC+Para. Values are displayed as means ± SEM. Only significant p values are shown.

Similar to HNSCC cells, DAC treatment in AML cell lines activated several aspects of COX-2-PGE_2_ pathway including upregulation of *PTGS2* and PGE_2_ receptors’ expression (Fig 7E). Upregulation of *CYP2E1* following DAC treatment was also noted for SKM-1 cells (Fig 7F). Ultimately, DAC increased ROS and mitochondrial superoxide production in these AML cells and this was further significantly enhanced by the addition of paracetamol but not valdecoxib (Fig 7G and H).

In further agreement with the results obtained for HNSCC cells, combined treatment did not add to the cell death caused by DAC alone as assessed by annexin V (Fig 7I). There was also little change to cell cycle progression after DAC when compared to control while combined treatment increased the number of cells retained in S phase when compared to DAC alone (Fig 7J). Following DAC and DAC-paracetamol treatments, AML cells displayed an increase of myeloid differentiation marker *CD11b* (*ITGAM*) (Fig 7K). Although this was not significantly different between combined treatment and DAC alone the effects of DAC+paracetamol were more apparent when cell morphology was assessed and showed a change towards myelocyte with increased cytoplasm and a high number of vacuoles (Fig 7L). These data indicate that DAC-paracetamol synergy and the effect it has on oxidative stress could be applicable to blood malignancies, where DAC is an accepted and widely used therapeutic option.

## Discussion

The search for new drugs to improve the poor survival rate of HNSCC is ongoing. Our preliminary results using four HNSCC cell lines and an *in vivo* mouse model show that DAC alone has therapeutic potential in HNSCC. However, the response is variable, as has been observed for other solid tumours [8] and patient stratification will be necessary to identify the DAC-responders. Future studies will look for predictive biomarkers, however the results shown here are clear that sensitivity to DAC is primarily dependent upon the ability of DAC to demethylate the DNA, and therefore likely due to incorporation, activation or retention of the drug, as suggested previously [39, 40]. Furthermore, an initial response to DAC treatment was a prerequisite for synergy with paracetamol, therefore the co-treatment is dependent upon the DNA demethylating capacity of DAC.

This is the first study to identify that paracetamol can enhance the anti-tumour activity of DAC. There are two main translational impacts of the DAC-paracetamol combination. Our data strongly indicate that adding paracetamol to DAC treatment could significantly lower the DAC dose needed to achieve therapeutic effects, broadening DAC application. In addition, it opens a potential for DAC dose reduction, potentially reducing DNA damage-related side effects.

Paracetamol is routinely prescribed as an analgesic, however so far it has not been considered whether its use may influence, positively or negatively, the efficacy of chemotherapy regimens. This study highlights that uncontrolled use of paracetamol during cancer therapies and clinical trials could affect the outcomes and interpretation of the results. Hence, further studies are required to look at the impact of supportive care medication in oncology.

Considering both DAC and paracetamol have broad effects with many potential interactions, we investigated and identified key mechanisms within the AA metabolism pathway that could underlie the synergy (summarized in Fig 5A). Both our data and analysis of public databases point towards DAC explicitly upregulating COX-2-PGE_2_ pathway, therefore providing cancer cells with survival advantage. This also reflects the previously reported bias in cancer cells towards PGE_2_ production to the detriment of other prostaglandins [41, 42]. It remains to be established whether the COX-2-PGE_2_ pathway activation is a direct result of DNA demethylation or is rather due to indirect mechanisms (e.g. response to dsRNA, cytokines or growth factors).

Surprisingly, the synergistic effect observed for paracetamol could not be reproduced using other COX-2 inhibitors. The data indicate an alternative paracetamol-specific mechanism; DAC-induced mimicry of paracetamol overdose leading to GSH depletion and exacerbated oxidative stress, both of which have the potential to restrict tumour growth and improve patient survival [37]. These cytoprotective pathways are also involved in development of drug resistance, hence their suppression might account for the extended drug sensitivity observed in AML cells after combined treatment.

Response to DAC has a profound effect on the transcriptional programme in HNSCC cells, the response is dominated by activation of type I interferon and anti-viral pathways, agreeing with recent reports on the role of ‘viral mimicry’ in cancer treatment with demethylating agents [3, 4]. These effects are maintained but not increased by combined treatment. Instead, the addition of paracetamol to DAC treatment led to a decrease in DNA, RNA and protein metabolism, together with reduced proliferation and enhanced differentiation. Concurrently, a strong enrichment for ‘keratinocyte differentiation’ was noted in the DSEA dataset shared by DAC and paracetamol signatures. Further experiments are required to establish the exact mechanisms leading to the changes in cell proliferation and differentiation upon combined treatment and whether they are linked to alterations in COX-2-PGE_2_ pathway and/or to ‘mimicry of paracetamol overdose’. One possibility involves the effects of AA metabolism and ROS on PI3K/Akt and MAPK pathways, the main proliferation drivers in HNSCC [43]. In HNSCC, mutations in the AA pathway have been reported to downregulate PI3K/Akt signalling [43], while patients with activating *PIK3CA* mutations specifically benefited from regular use of NSAIDs [44]; confirming the significance of AA and COX pathways in HNSCC development and treatment.

This study provides evidence that DAC may have anti-tumour potential in a subset of HNSCC cases, highlighting the need for effective biomarkers and patient stratification. We describe how this response can be enhanced by the presence of paracetamol, applied in doses commonly used to alleviate pain. Synergy was observed in AML and DAC-responsive HNSCC cells, with similar genes and pathways affected in both. The clinical implications of this are twofold. Firstly, the addition of paracetamol could allow for DAC dose reduction, reducing DNA damage-related side effects and widening clinical usability, possibly to solid tumours, including HNSCC. Secondly, we highlight the fact that the commonly used drug, available as over-the-counter medicine and often self-medicated by patients, can change the cancer cell response to a chemotherapeutic. Therefore, considering the mechanisms described here, paracetamols interaction with other drugs, especially chemotherapeutics, should be taken into consideration in cancer treatment. This manuscript provides a solid rationale for the controlled use of paracetamol in AML, where DAC treatment already has been approved, and suggests efficacy may also be applicable to HNSCC. Since paracetamol is a very cheap and relatively safe drug, it could be added to treatment with minimal cost but considerable translational impact.

## Materials and methods

### Cell lines and culture conditions

Five human HNSCC cell lines were used: SCC040 (German Culture Collection, DSMZ (#ACC660)), FaDu (ATCC (HTB-43)), VU40T (Prof H. Joenje (VU University Medical Centre, Amsterdam)), HN12 (Dr J.F. Ensley (Wayne State University, Detroit, MI)) and UDSCC2 (Dr Henning Bier (University of Duesseldorf, Germany)). The cell lines were authenticated using STR profiling (NorthGene, UK). All HNSCC cell lines were maintained in DMEM (Sigma-Aldrich) supplemented with 10% fetal bovine serum (FBS) (Sigma-Aldrich), 1% penicillin-streptomycin (Gibco), 4 mM L-glutamine (Sigma-Aldrich), 1X non-essential amino acids (Life Technologies) and 1 mM sodium pyruvate (Life Technologies).

Primary human oral keratinocyte (HOK) cells were purchased from Caltag Medsystems and cultured over Poly-L-Lysine in Oral Keratinocyte Medium supplemented with 1% oral keratinocyte growth supplement (all purchased from ScienceCell) and 1% penicillin-streptomycin.

Two AML cell lines were used: SKM-1 (from Dr Stefan Heinrichs, University of Essen, Germany) and HL-60 (ATCC (CCL-240)). Cells were cultured in RPMI1640 medium (ThermoFisher Scientific), supplemented with 15% FBS (Sigma-Aldrich), 1% L-glutamine and 1% penicillin-streptomycin.

All cell lines were regularly tested for mycoplasma using MycoAlert (Lonza) and experiments were performed within 18 passages after thawing.

### Drug treatments

All drugs were purchased from Sigma-Aldrich. 5-Aza-2’-deoxycytidine (DAC) was dissolved in either 50% acetic acid or in ≥99.9% dimethyl sulfoxide (DMSO), all other drugs were dissolved in DMSO. The treatments were carried out for 96h (HNSCC cells) or 72h (AML cells) using a relevant vehicle as control.

### Viability assay

Relative viability was determined in 96 well plates using the CellTiter-Blue Cell Viability Assay (Promega), including a minimum of triplicate wells per sample, vehicle, high concentration vehicle and ‘media only’ controls. Background absorbance was accounted for by subtracting the media only control from each sample and then these were normalised to a vehicle-only control. Sigma plot software (Systat Software Inc.) was used to generate sigmoidal, 4-parameter dose response curves after drug titrations. IC_50_ values were calculated using MyCurveFit (MyAssays Ltd.).

### DAC sensitising assay

A panel of 100 drugs (Drug Library FMC1) were administered at the reported peak serum concentrations (Cmax) [13]. Cells were seeded in 96-well plates and treated with each drug alone or in combination with 500 nM DAC. The assay was performed blind in biological triplicates with controls hidden within the panel.

### Determining synergy

The Chou-Talalay method was employed to determine synergy [25] using constant-ratio matched titrations (0.125X, 0.25X, 0.5X, 1X, 2X, 3X, 4X, and 8X) of multiples of the Cmax (500 nM for DAC, 132 μM for paracetamol) and assessed by cell viability. The CompuSyn software [25] was used to calculate combination index (CI) and dose reduction index (DRI) values. CI values below 1 indicate synergy. DRI values indicate how many times less each drug can be used when in combination.

### Long treatment of AML cells and growth rate assessment

SKM-1 and HL-60 cells were subjected to a repeated cycle of treatments. The cells were plated in 6-well plates at 0.5×10^6^ cells/ml and 0.375×10^6^ cells/ml for SKM-1 and HL-60, respectively, treated for 72h after which the cells were counted and re-plated without drugs at original concentrations. The withdrawal period lasted 21 days during which the cells were passaged twice a week. The treatment cycle was repeated four times and the cells were counted at each passage using trypan blue (Sigma-Aldrich) staining and haemocytometer cell counting. Growth rate was calculated using the following equation Gr=ln(N(t)/N(0))/t, where Gr = growth rate, N(t)=number of cells at time t, N(0)=number of cells at time 0, and t=time(hours).

### DNA dot blotting

DNA was extracted using the DNeasy Blood and Tissue Kit (Qiagen) as described by the manufacturer. Dot blotting was performed as described in Current Protocols in Molecular Biology using a titration of DNA (1). Dried membranes (Nytran Supercharge Positively Charged Nylon Membrane (Fisher Scientific)) were blocked in 5% low-fat milk in PBS for 1h and incubated with primary antibody (anti-5mC, Cell Signalling (28692S), 1:750 dilution) in 1% bovine serum albumin (BSA) at 4°C overnight. Membranes were incubated with secondary antibody (Santa Cruz, Cat# sc-2004, 1:5000 dilution) for 2h at 4°C. Membranes were developed using 1:1 Amersham ECL western blotting reagent (GE Healthcare) to SuperSignal West Femto Chemiluminescent Substrate (Thermo Fisher Scientific) and detected by autoradiography. Blots were stained for total DNA by 30min incubation in 0.04% methylene blue (Sigma-Aldrich). ImageJ software (2) was used to quantify the intensity of the 1 μg and 0.5 μg dots and each 5-methylcytosine dot was normalised against its methylene blue counterpart. For each biological replicate an average was taken from the normalised 1 μg and 0.5 μg dots and the technical replicates for use in further analysis.

### Western Blot analysis

Cell pellets were incubated with RIPA buffer (150mM sodium chloride, 0.5% sodium deoxycholate, 0.1% sodium dodecyl sulphate, 50mM Tris-HCl pH8) supplemented with protease inhibitors (Roche) for 45min then centrifuged at 17,000g for 10min and the supernatant was retained. Proteins were denatured in Laemmli buffer (4% SDS, 20% glycerol, 10% 2-mercaptoethanol, 0.004% bromophenol blue and 0.125 M Tris HCl), heated to 95°C for 5min and separated on a 10% SDS-PAGE gel before transferring onto a nitrocellulose membrane (Amersham Hybond ECL, GE Healthcare) using BioRad transfer apparatus. The protein transfer was confirmed with Ponceau S staining (SigmaAldrich). Membranes were blocked in 20% milk in PBST for 1h following primary antibody incubation (COX-2: Abcam, Cat# ab15191, 1:1000 dilution, 1h at room temperature in 1% BSA; CYP2E1: Abcam, Cat# ab28146, 1:2500 dilution, overnight 4oC in 5% milk; lamin A/C: Santa Cruz, Cat# sc-20681, 1:10000, 1h at room temperature in 5% milk). Membranes were incubated with secondary antibody (anti-rabbit-IgG-HRP Santa Cruz, Cat# sc-2004, 1:5000 dilution) for 2h and developed as described for DNA dot blotting. ImageJ software was used to quantify the intensity of each band and bands of interest were normalised against the corresponding lamin A/C bands.

### Giemsa-Jenner staining

VU40T cells were grown on coverslips for 96h, fixed in methanol and stained with Giemsa and Jenner (VWR) as previously described [13]. SKM-1 cells were transferred to a glass slide with a Cytospin 3 (Thermo Shandon) before fixation. Microscope images were taken using EVOS XL Core Imaging System or BX-50 Olympus with a 100x oil immersion lens.

### Immunofluorescence analysis

Immunofluorescence analysis was undertaken as described in (Roulois et al, 2015). For γH2AX staining, cells were first incubated with ice-cold pre-extraction buffer (10 mM PIPES pH 6.8, 300 mM sucrose, 20 mM NaCl, 3 mM MgCl2 0.5% Triton X-100) for 7min. For both γH2AX and Ki67 staining the cells were fixed in methanol for 15min, followed by 1min incubation with ice-cold acetate. Samples were blocked with 1% BSA in PBS for 1h at 4***°***C following incubation with the primary antibodies overnight (Ki67 (Abcam, Cat# ab15580) 1:1000; γH2AX (phospho S139) (Abcam, Cat# ab2893) 1:1000). Coverslips were incubated with secondary antibody (Jackson ImmunoResearch, Donkey anti-rabbit Alexa Fluor 488 cat# 711-545-152, 1:250) for 1h at room temperature, washed with PBS before drying and mounting using ProLong Gold Antifade Mountant with DAPI (Fisher Scientific). Cells were stored in the dark for 3 days prior to imaging on a Zeiss 780 Zen confocal microscope. For Ki67 staining, ImageJ was used to count percentage of positive cells in triplicate images from each biological replicate. For γH2AX staining, the number of γH2AX per nuclei was counted manually across all focal planes. 100 nuclei were analysed in each biological replicate.

### Apoptosis assay and cell cycle analysis

Apoptosis and necrosis were assessed using the Annexin V Apoptosis Detection APC kit (eBioscience, ThermoFisher Scientific) as the manufacturer recommends. For cell cycle analysis, cells were fixed in 70% ice cold ethanol and resuspended in 200 μl 50 μg/ml propidium iodide solution (Sigma-Aldrich) and 50 μl of 100 μg/ml RNase solution (Roche). The solutions were analysed by flow cytometry on a Cyan B FACS analyser (Beckman Coulter).

Following flow cytometry, the cells were gated into four populations and counted: healthy cells which were positive for neither PI nor annexin V; necrotic cells stained only with PI; early apoptotic cells stained only with annexin V and late apoptotic cells stained with both PI and annexin V. The proportion of cells in each quadrant after 48h, 72h, 96h of treatment was counted and compared to the vehicle control.

### ELISA

The levels of PGE_2_, Leukotriene B_4_ (LTB_4_) and Cysteinyl leukotrienes (LTC_4_, LTD_4_, and LTE_4_) were assessed in cell media using respective ELISA kits (Abcam). The results were normalised to the corresponding CellTiter-Blue viability results or cell count.

### Determining glutathione concentration

The glutathione levels were measured using the GSH-Glo Glutathione Assay (Promega) with 1 mM paracetamol used as a positive control. The results were normalized against the corresponding CellTiter-Blue cell viability.

### NAC rescue assay

N-acely-l-cysteine (NAC, Sigma-Aldrich) was dissolved in water and neutralised using 1 M sodium hydroxide (NaOH) (Sigma-Aldrich). VU40T cells were treated with 2.5 mM NAC or an equivalent volume of vehicle for 48h. Following this, wells were washed with fresh media and cells were treated with DAC, paracetamol or both for 96h and cell viability was determined.

### Assessment of reactive oxygen species (ROS) and mitochondrial superoxide (mitosox)

MitoSOX Red (Molecular Probes, Cat# M36008, ThermoFisher Scientific) was used to assess mitosox. PBS-washed cells were resuspended in 200 μl PBS (37°C) containing 5 μM MitoSOX Red, incubated at 37°C for 10 min and then analysed by flow cytometry (emission wavelength 580 nm). Reactive oxygen species (ROS) were measured using carboxy-H2DCFDA (5-(and-6)-carboxy-2’,7’-dichlorodihydrofluorescein diacetate, Cat# C369, Invitrogen). 10 μM carboxy-H2DCFDA was added to 0.5 ml cell suspension for 45 min, cells were washed with PBS and resuspended in 200 μl PBS (37°C). Resuspended cells were analysed by flow cytometry (emission wavelength 517-527 nm) with BD FACS Calibur and BD Cell Quest software. The geometric mean was calculated for each sample and experimental condition.

### Real-Time Quantitative PCR (qRT-PCR)

Total RNA was extracted using RNeasy Mini Kit (Qiagen), including DNase I digestion step (RNase-Free DNase set, Qiagen). 1 μg of RNA was reverse transcribed (iScript cDNA Synthesis Kit (BioRad)) and the cDNA was purified with QiaQuick PCR Purification Kit (Qiagen). Purified cDNA was quantified using Qubit dsDNA High Sensitivity Assay kit (Thermo Fisher Scientific) to support normalization. qRT-PCRs were performed on a LightCycler 480 II using LightCycler 480 SYBR Green (Roche).

The primers used in qRT-PCR are listed in Supplementary Table S7. Relative RNA values were normalised against the concentration of the purified cDNA as measured by Qubit dsDNA High Sensitivity Assay kit (Thermo Fisher Scientific). This method of normalization was treated as an additional step because the DAC treatments can lead to decrease of RNA yield and changes in housekeeping gene expression. Each sample was examined at least in triplicate. PCR product specificity was confirmed by a melting-curve analysis. For analysis of qRT-PCR results the PRISM software was used to generate bar graphs and calculate parametric paired t-tests comparing treated samples to vehicle only, and combined treatments to individual drugs.

### RNA sequencing

For each sample three biological RNA replicates were pooled to make a library using the TruSeq Stranded mRNA Library Prep kit and sequenced using Illumina NextSeq 500 in paired-end mode at 2×75 bases in the Genomics Birmingham facility (Birmingham, UK). Reads were aligned to the genome (hg19) using HiSAT2 and processed with bedtools to generate normalised coverage plots.

Quantification of the RNA sequencing data was performed according to the latest recommended pipeline as defined in the DeSeq2 software. A count for each gene was calculated using the reference-free aligner Salmon (Patro et al, 2017) and the resulting count table was processed using DeSeq2 (Love et al, 2014) to compare treatment groups. There were 1200 upregulated (log2 fold change ≥1) and 937 downregulated (log2 fold change ≤-1) genes, further divided into six groups: i) genes up-regulated by paracetamol (n=151); ii) genes down-regulated by paracetamol (n=127); iii) genes up-regulated by DAC (n=658); iv) genes down-regulated by DAC (n=293); v) genes up-regulated by combined treatment (n=915); and vi) genes down-regulated by combined treatment (n=721). Each group was subjected to gene ontology (GO) analysis using the Gene Ontology Consortium software (Ashburner et al, 2000) followed by removal of redundant terms using REVIGO (Supek et al, 2011). The top 5 terms with the Log10 p-value ≤3.0 from each group were collated; none of the GO terms from the gene set downregulated by paracetamol fulfilled this condition. The Log10 p-values were used to generate heatmap.

### Mouse xenograft study

All animal studies were performed in accordance with the UK Home Office Animal (Scientific Procedures) Act 1986 and approved by the local University of Birmingham Ethical Review Committee.

The mouse xenograft study was performed as described previously [38] and is described briefly below.

To examine the toxicity and anti-tumour efficacy of DAC and paracetamol, we utilised male NOD/SCID/gamma (NSG) mice (Charles River) which were kept in 12 hour light and 12 hour dark cycles in individually ventilated cages. Mice were maintained on mouse feed and water, ad libitum, and were at least 6 weeks old at the start of treatment.

Toxicity studies with 3 animals per single treatment and combination treatment were used to identify safe doses for use in subsequent efficacy study: 0.4 mg/kg DAC and 100 mg/kg paracetamol. In the toxicity studies the drugs were given 5 days a week for three weeks, first for DAC and paracetamol separately and then in combination.

To assess the efficacy, 24 male NSG mice were implanted with 5×10^6^ FaDu HNSCC cells suspended in serum-free medium and injected subcutaneously into the right flank. Tumours were given three days to become established, at which point mice were randomly allocated into four treatment groups, each group containing 6 animals, as follows: (1) 0.4 mg/kg DAC dissolved in PBS via intraperitoneal (IP) injection on a 5 day on, 2 day off regimen from day 4 onwards; (2) 100 mg/kg paracetamol dissolved in PBS given via oral gavage following the same regimen; (3) DAC plus paracetamol as above; (4) control (PBS) given both through the oral gavage and IP injection. The administration routes were chosen based on previously published literature: IP administration of DAC has previously been described in [45] while paracetamol is commonly administered intraorally [46, 47]. Animals were monitored daily for signs of ill health and the tumours were measured. Tumour measurements were taken using callipers and volumes calculated using the formula L x W^2^. Mice were culled once tumours reached a maximum of 1250 mm^3^; tumours became ulcerated; when animals showed signs of ill health; or at the completion of the study. One mouse from group 1 had to be excluded on Day 10 due to dosing error. When displaying tumour growth over time, to gain an understanding of efficacy over a longer period of time, data were plotted until at least 4 animals per group remained; when fewer animals remained in a group, no more data were plotted. The tumour growth is also shown as time to reach volume of 200 mm^3^; tumours smaller than 200 mm^3^ at the end of the study (day 25) were counted as ‘Day 25’. The difference in tumour size for each group was shown at day 15 (Control n=6, paracetamol n=6, DAC n=5, DAC+paracetamol n=6) and at the end of the study (day 25: DAC n=3 and DAC+paracetamol n=5).

Tumours were excised and snap frozen for subsequent analysis. Tumour tissue was available from 6 animals in control and paracetamol-treated groups, 4 animals from DAC-treated and 5 animals from DAC+paracetamol treated groups. Approximately 10-20mg tissue was pulverized and total RNA extracted as described before (see: Real-time quantitative PCR).

### Quantification and statistical analysis

Statistical analysis and graphs were performed using GraphPad PRISM 8 software, unless stated otherwise. Details of the statistical tests used for each experiment are given in the corresponding figure legends.

### TCGA data

The following data were retrieved from The Cancer Genome Atlas (TCGA) via cBioPortal for Cancer Genomics (Cerami et al, 2012; Gao et al, 2013): (i) expression and genetic alterations in *TP63* gene available for 496 patients (Appendix Fig.S2A); (ii) percentage of samples for each cancer type with mRNA Expression z-score threshold ± 2.0 (RNA Seq V2 RSEM) for all cancers with available TCGA provisional data (Appendix Tables S5 and S6); (iii) clustered gene expression heatmap for genes in COX-2 pathway in HNSCC tumours (520 patients, Fig. 3F); (iv) Logrank test p-values for overall survival and/or disease/progression-free survival for all cancer with available TCGA provisional data in indicated set of genes (expression z-score threshold ± 2.0), (Appendix Tables S5 and S6, Fig. 5B-D, Appendix Fig. S6); (v) Kaplan-Meier curves for HNSCC (520 patients) disease-progression free survival based on gene expression alterations in indicated sets of genes. cBioPortal was used to determine patients with increased expression of the genes indicated. The total number of HNSCC cases available were then split depending on whether they had one or more of these genes upregulated (red lines) or none (blue lines). The total number of cases within each group is shown as an n number on the graphs. The following gene sets have been used: COX-2 pathway (*PTGS2, PTGES, PTGES2, PTGES3, PTGER1, PTGER2, PTGER3, PTGER4*; Fig. 5B); Glutathione synthesis (*GSS, GCLC, GCLM, GGCT, OPLAH, GSR*; Fig. 5C); Antioxidant response genes down-regulated by DAC (*PRDX1, PRDX6, SRXN1, UGT1A1, NQO1, FTL, AKR1C2*; Fig. 5D); Thioredoxin pathway genes down-regulated by DAC (*TXN2, SLC7A11, TXN, TXNRD1*; Fig. 5D); COX-2-PGF2, -PGI2, - PGD2, -TBX2 pathways (*PTGS2, PTGDR2, PRXL2B, PTGDS, PTGIS, PTGIR, PTGFR, TBXA2R, TBXAS1*; Appendix Fig. S6A); LOX pathway (ALOX5, ALOX15, ALOX15B, ALOX12, LTA4H; Appendix Fig. S6B).

### Drug perturbation signatures

Drug perturbation signatures were downloaded for the BROAD Connectivity Map dataset (CMAP) using the PharmacoGx package (version 1.14.0) (Smirnov et al, 2016) in R. cMap drug perturbation signatures represent transcriptionally profiled, drugs-treated cancer cell lines involving 11833 genes and 1309 drugs across 5 cancer cell lines (breast cancer MCF7 and ssMCF7, pancreatic cancer PC3, melanoma SKMEL5 and AML HL-60). Precomputed signatures for CMAP were available for 1288 drugs; the signatures are calculated using a linear regression model adjusted for treatment duration, cell line identity, and batch, as described previously (Smirnov et al., 2016). Heatmaps of drug perturbation signatures (transcriptional profiles) for Decitabine (DAC), paracetamol (Para), and Valdecoxib were plotted using ggplots package (version 3.0.1.1).

All analyses have been conducted using the R statistical software (version 3.6.0); listed software dependencies are available on the Comprehensive Repository R Archive Network (CRAN) or Bioconductor (BioC).

### Drug Set Enrichment Analysis

The effect of Decitabine, paracetamol and valdecoxib drug combinations on KEGG pathways and Gene Ontology Biological Processes was conducted using the Drug Set Enrichment Analysis (DSEA) server [36]. Significance of the perturbed pathways of interest was identified using the corresponding p-values per geneset; log_10_ (p-values) were plotted as heatmaps.

## Data availability

RNA-Seq data: The data are deposited at the GEO repository, accession number GSE110045 and SRA, accession number SRP132039.

## Acknowledgments

We would like to thank Dr Jo Parish (University of Birmingham, UK) for critically reviewing the manuscript. This research was supported by grants from the FP7 Framework Marie Curie Actions CIG (methDRE) to M. Wiench and The Royal Society (RS-9-10-2012) to M. Wiench. H.J. Gleneadie received a joint University of Birmingham and Medical Research Council fellowship.

## Author contributions

H.J.G., M.W., F.L.K., B.A.S., and P.R.C designed the project. H.J.G., M.W., and F.L.K. analysed the data. H.J.G. and M.W. performed most of the experiments in HNSCC cell lines. A.B. and P.G. performed most of the experiments in AML cell lines. N.B. designed and analysed drug synergy experiments. J.B. and N.B. performed *in vivo* experiments. Y.J. performed and analysed ROS and mitosox assays. S.C. analysed RNA-seq data. D.M.A.G. performed DSEA analysis. S.R., J.S.G. and A.M. provided essential advice and material. M.W. and H.J.G. wrote the manuscript with input from S.R., F.L.K., S.C. and H.M.

## References

1 Jones PA, Baylin SB. The fundamental role of epigenetic events in cancer. Nat Rev Genet 2002; 3: 415–428.

2 Stresemann C, Lyko F. Modes of action of the DNA methyltransferase inhibitors azacytidine and decitabine. Int J Cancer 2008; 123: 8–13.

3 Chiappinelli KB, Strissel PL, Desrichard A, Li H, Henke C, Akman B et al. Inhibiting DNA Methylation Causes an Interferon Response in Cancer via dsRNA Including Endogenous Retroviruses. Cell 2015; 162: 974–986.

4 Roulois D, Loo Yau H, Singhania R, Wang Y, Danesh A, Shen SY et al. DNA-Demethylating Agents Target Colorectal Cancer Cells by Inducing Viral Mimicry by Endogenous Transcripts. Cell 2015; 162: 961–973.

5 Kantarjian HM, Thomas XG, Dmoszynska A, Wierzbowska A, Mazur G, Mayer J et al. Multicenter, randomized, open-label, phase III trial of decitabine versus patient choice, with physician advice, of either supportive care or low-dose cytarabine for the treatment of older patients with newly diagnosed acute myeloid leukemia. J Clin Oncol 2012; 30: 2670–2677.

6 Nieto M, Demolis P, Behanzin E, Moreau A, Hudson I, Flores B et al. The European Medicines Agency Review of Decitabine (Dacogen) for the Treatment of Adult Patients With Acute Myeloid Leukemia: Summary of the Scientific Assessment of the Committee for Medicinal Products for Human Use. Oncologist 2016; 21: 692–700.

7 Tsai HC, Li H, Van Neste L, Cai Y, Robert C, Rassool FV et al. Transient low doses of DNA-demethylating agents exert durable antitumor effects on hematological and epithelial tumor cells. Cancer Cell 2012; 21: 430–446.

8 Linnekamp JF, Butter R, Spijker R, Medema JP, van Laarhoven HWM. Clinical and biological effects of demethylating agents on solid tumours - A systematic review. Cancer Treat Rev 2017; 54: 10–23.

9 Simard EP, Torre LA, Jemal A. International trends in head and neck cancer incidence rates: differences by country, sex and anatomic site. Oral Oncol 2014; 50: 387–403.

10 Squier CA, Kremer MJ. Biology of oral mucosa and esophagus. J Natl Cancer Inst Monogr 2001: 7–15.

11 Steinmann K, Sandner A, Schagdarsurengin U, Dammann RH. Frequent promoter hypermethylation of tumor-related genes in head and neck squamous cell carcinoma. Oncol Rep 2009; 22: 1519–1526.

12 Abele R, Clavel M, Dodion P, Bruntsch U, Gundersen S, Smyth J et al. The EORTC Early Clinical Trials Cooperative Group experience with 5-aza-2’-deoxycytidine (NSC 127716) in patients with colo-rectal, head and neck, renal carcinomas and malignant melanomas. Eur J Cancer Clin Oncol 1987; 23: 1921–1924.

13 Khanim FL, Merrick BA, Giles HV, Jankute M, Jackson JB, Giles LJ et al. Redeployment-based drug screening identifies the anti-helminthic niclosamide as anti-myeloma therapy that also reduces free light chain production. Blood Cancer J 2011; 1: e39.

14 Bertolini A, Ferrari A, Ottani A, Guerzoni S, Tacchi R, Leone S. Paracetamol: new vistas of an old drug. CNS Drug Rev 2006; 12: 250–275.

15 Graham GG, Davies MJ, Day RO, Mohamudally A, Scott KF. The modern pharmacology of paracetamol: therapeutic actions, mechanism of action, metabolism, toxicity and recent pharmacological findings. Inflammopharmacology 2013; 21: 201–232.

16 Wang D, Dubois RN. Eicosanoids and cancer. Nat Rev Cancer 2010; 10: 181–193.

17 Liu B, Qu L, Yan S. Cyclooxygenase-2 promotes tumor growth and suppresses tumor immunity. Cancer Cell Int 2015; 15: 106.

18 Saba NF, Choi M, Muller S, Shin HJ, Tighiouart M, Papadimitrakopoulou VA et al. Role of cyclooxygenase-2 in tumor progression and survival of head and neck squamous cell carcinoma. Cancer Prev Res (Phila) 2009; 2: 823–829.

19 Kurtova AV, Xiao J, Mo Q, Pazhanisamy S, Krasnow R, Lerner SP et al. Blocking PGE2-induced tumour repopulation abrogates bladder cancer chemoresistance. Nature 2015; 517: 209–213.

20 Kim YY, Lee EJ, Kim YK, Kim SM, Park JY, Myoung H et al. Anti-cancer effects of celecoxib in head and neck carcinoma. Mol Cells 2010; 29: 185–194.

21 Kobrinsky NL, Hartfield D, Horner H, Maksymiuk A, Minuk GY, White DF et al. Treatment of advanced malignancies with high-dose acetaminophen and N-acetylcysteine rescue. Cancer Invest 1996; 14: 202–210.

22 Posadas I, Vellecco V, Santos P, Prieto-Lloret J, Cena V. Acetaminophen potentiates staurosporine-induced death in a human neuroblastoma cell line. Br J Pharmacol 2007; 150: 577–585.

23 Wu YJ, Neuwelt AJ, Muldoon LL, Neuwelt EA. Acetaminophen enhances cisplatin- and paclitaxel-mediated cytotoxicity to SKOV3 human ovarian carcinoma. Anticancer Res 2013; 33: 2391–2400.

24 European Medicines Agency E. Dacogen. In: Agency EM (ed), vol. 2019: https://www.ema.europa.eu/en/documents/product-information/dacogen-epar-product-information_en.pdf, 2019.

25 Chou TC. Theoretical basis, experimental design, and computerized simulation of synergism and antagonism in drug combination studies. Pharmacol Rev 2006; 58: 621–681.

26 Yoh K, Prywes R. Pathway Regulation of p63, a Director of Epithelial Cell Fate. Front Endocrinol (Lausanne) 2015; 6: 51.

27 Ashburner M, Ball CA, Blake JA, Botstein D, Butler H, Cherry JM et al. Gene ontology: tool for the unification of biology. The Gene Ontology Consortium. Nat Genet 2000; 25: 25–29.

28 Supek F, Bosnjak M, Skunca N, Smuc T. REVIGO summarizes and visualizes long lists of gene ontology terms. PLoS One 2011; 6: e21800.

29 Park SW, Heo DS, Sung MW. The shunting of arachidonic acid metabolism to 5-lipoxygenase and cytochrome p450 epoxygenase antagonizes the anti-cancer effect of cyclooxygenase-2 inhibition in head and neck cancer cells. Cell Oncol (Dordr) 2012; 35: 1–8.

30 Lamb J, Crawford ED, Peck D, Modell JW, Blat IC, Wrobel MJ et al. The Connectivity Map: using sene-expression signatures to connect small molecules, genes, and disease. Science 2006; 313: 1929–1935.

31 Dahlin DC, Miwa GT, Lu AY, Nelson SD. N-acetyl-p-benzoquinone imine: a cytochrome P-450-mediated oxidation product of acetaminophen. Proc Natl Acad Sci U S A 1984; 81: 1327–1331.

32 Forman HJ, Zhang H, Rinna A. Glutathione: overview of its protective roles, measurement, and biosynthesis. Mol Aspects Med 2009; 30: 1–12.

33 Ray PD, Huang BW, Tsuji Y. Reactive oxygen species (ROS) homeostasis and redox regulation in cellular signaling. Cell Signal 2012; 24: 981–990.

34 Raghunath A, Sundarraj K, Nagarajan R, Arfuso F, Bian J, Kumar AP et al. Antioxidant response elements: Discovery, classes, regulation and potential applications. Redox Biol 2018; 17: 297–314.

35 Telorack M, Meyer M, Ingold I, Conrad M, Bloch W, Werner S. A Glutathione-Nrf2-Thioredoxin Cross-Talk Ensures Keratinocyte Survival and Efficient Wound Repair. PLoS Genet 2016; 12: e1005800.

36 Napolitano F, Sirci F, Carrella D, di Bernardo D. Drug-set enrichment analysis: a novel tool to investigate drug mode of action. Bioinformatics 2016; 32: 235–241.

37 Traverso N, Ricciarelli R, Nitti M, Marengo B, Furfaro AL, Pronzato MA et al. Role of glutathione in cancer progression and chemoresistance. Oxid Med Cell Longev 2013; 2013: 972913.

38 Bryant J, Batis N, Franke AC, Clancey G, Hartley M, Ryan G et al. Repurposed quinacrine synergizes with cisplatin, reducing the effective dose required for treatment of head and neck squamous cell carcinoma. Oncotarget 2019; 10: 5229–5244.

39 Qin T, Jelinek J, Si J, Shu J, Issa JP. Mechanisms of resistance to 5-aza-2’-deoxycytidine in human cancer cell lines. Blood 2009; 113: 659–667.

40 Wu L, Shi W, Li X, Chang C, Xu F, He Q et al. High expression of the human equilibrative nucleoside transporter 1 gene predicts a good response to decitabine in patients with myelodysplastic syndrome. J Transl Med 2016; 14: 66.

41 Cebola I, Custodio J, Munoz M, Diez-Villanueva A, Pare L, Prieto P et al. Epigenetics override pro-inflammatory PTGS transcriptomic signature towards selective hyperactivation of PGE2 in colorectal cancer. Clin Epigenetics 2015; 7: 74.

42 Cebola I, Peinado MA. Epigenetic deregulation of the COX pathway in cancer. Prog Lipid Res 2012; 51: 301–313.

43 Cancer Genome Atlas N. Comprehensive genomic characterization of head and neck squamous cell carcinomas. Nature 2015; 517: 576–582.

44 Hedberg ML, Peyser ND, Bauman JE, Gooding WE, Li H, Bhola NE et al. Use of nonsteroidal anti-inflammatory drugs predicts improved patient survival for PIK3CA-altered head and neck cancer. J Exp Med 2019; 216: 419–427.

45 Ecke I, Petry F, Rosenberger A, Tauber S, Monkemeyer S, Hess I et al. Antitumor effects of a combined 5-aza-2’deoxycytidine and valproic acid treatment on rhabdomyosarcoma and medulloblastoma in Ptch mutant mice. Cancer Res 2009; 69: 887–895.

46 Takehara M, Hoshino T, Namba T, Yamakawa N, Mizushima T. Acetaminophen-induced differentiation of human breast cancer stem cells and inhibition of tumor xenograft growth in mice. Biochem Pharmacol 2011; 81: 1124–1135.

47 Skoglund LA, Ingebrigtsen K, Lausund P, Nafstad I. Plasma concentration of paracetamol and its major metabolites after p.o. dosing with paracetamol or concurrent administration of paracetamol and its N-acetyl-DL-methionine ester in mice. Gen Pharmacol 1992; 23: 155–158.

