## Supplemental info for "The common analgesic paracetamol enhances the anti-tumour activity of decitabine through exacerbation of oxidative stress"

The Appendix contains the following supplementary information:

SI Tables S1 to S7 (Tables S3 and S4 as separate .xlsx files)

SI Figures S1 to S7

### SI Supplementary Tables

Table S1. DRI (Dose Reduction Index) for VU40T, combined treatment.

| Fa (%) | VU40T | | | |
|---|---|---|---|---|
| | Dose | | DRI | |
| | DAC | Para | DAC | Para |
| **10** | 0.01 | 105 | 0.30 | 10.18 |
| **25** | 0.16 | 251 | 1.22 | 7.11 |
| **50** | 2.26 | 595 | **4.99** | **4.97** |
| **75** | 31 | 1414 | 20.35 | 3.47 |
| **97** | 9300 | 9200 | 426 | 1.59 |

Table S2. DRI for HN12, combined treatment.

| Fa (%) | HN12 | | | |
|---|---|---|---|---|
| | Dose | | DRI | |
| | DAC | Para | DAC | Para |
| **10** | 0.004 | 186 | 0.49 | 80 |
| **25** | 0.06 | 1962 | 1.19 | 147 |
| **50** | 0.85 | 20653 | **2.92** | **269** |
| **75** | 11.92 | 217412 | 7.15 | 493 |
| **97** | 3630 | 3.55E+07 | 49.5 | 1827 |

Table S3. Differentially expressed (increased and decreased) genes in VU40T cells following DAC, paracetamol and DAC+paracetamol treatments. See the Table_S3.xlsx file.

Table S4. Full list of GO terms from REVIGO for differentially expressed groups (DAC, paracetamol and DAC+paracetamol treatments). See the Table_S4.xlsx file.

Table S5. cBioPortal RNA-seq data (for cancers with provisional TCGA data): frequency of expression alterations (%) in genes from COX-2-$PGE_2$ pathway and correlation to survival (Logrank test p-value).

| **Cancer type** | **n** | **PTGS2** | **PTGES** | **PTGES2** | **PTGES3** | **PTGER1** | **PTGER2** | **PTGER3** | **PTGER4** | **OS** | **DFS** |
|---|---|---|---|---|---|---|---|---|---|---|---|
| Adrenocortical ca. | 79 | 1.3 | 4 | 5 | 19 | 9 | 1.3 | 5 | 6 | 0.181 | 0.130 |
| Cholangiocarcinoma | 36 | 6 | 6 | 6 | 2.8 | 8 | 2.8 | 2.8 | 2.8 | 0.322 | |
| Bladder urothelial ca. | 408 | 4 | 3 | 2.5 | 8 | 5 | 3 | 4 | 8 | 0.0702 | |
| Colorectal adenoca. | 379 | 3 | 6 | 1.6 | 7 | 2.4 | 4 | 4 | 4 | 0.0375 | |
| Breast invasive ca. | 1093 | 1.7 | 4 | 4 | 8 | 1.6 | 2.6 | 2.8 | 3 | 0.259 | |
| Glioblastoma mult. | 160 | 3 | 6 | 6 | 8 | 6 | 4 | 6 | 3 | | |
| Cervical SCC | 304 | 2.6 | 6 | 4 | 9 | 5 | 4 | 8 | 6 | 0.17* | |
| Esophageal carcinoma | 184 | 7 | 2.7 | 8 | 8 | 5 | 6 | 1.6 | 9 | | |
| Stomach adenoca. | 415 | 5 | 4 | 9 | 9 | 4 | 0.7 | 4 | 7 | | 0.09* |
| Uveal melanoma | 80 | 5 | 2.5 | 6 | 6** | 5 | 5 | 5 | 5 | | |
| HNSCC | 520 | 2.3 | 4 | 7 | 7 | 2.5 | 4 | 3 | 6 | 0.090 | 5.693e-3 |
| Kidney renal clear cell ca | 533 | 1.3 | 2.3 | 2.4 | 5 | 0.4 | 4 | 7 | 5 | 0.179 | 0.144 |
| Kidney renal papillary cell ca. | 290 | 2.4 | 4 | 4 | 9 | 3 | 4 | 2.4 | 4 | 0.034 | 0.307 |
| Liver hepatocellular ca. | 371 | 0.8 | 6 | 7 | 8 | 6 | 3 | 1.3 | 4 | 0.283 | |
| Lung adenocarcinoma | 515 | 4 | 4 | 3 | 11 | 2.1 | 5 | 3 | 4 | 0.039 | |
| Lung SCC | 501 | 4 | 6 | 3 | 5 | 2.4 | 1.2 | 1.4 | 7 | | 0.202 |
| AML | 173 | 4 | 5 | 4 | 2.9 | 2.3 | 3 | 2.9 | 2.3 | | n/a |
| Ovarian serous cystadenoca. | 307 | 0.7 | 2.6 | 7 | 4 | 2.3 | 2 | 5 | 5 | 2.305e-3 | 0.286 |
| Pancreatic adenoca. | 178 | 4 | 4 | 1.7 | 10 | 3 | 4 | 6 | 4 | 0.089 | 0.080 |
| Mesothelioma | 87 | 3 | 1.1 | 1.1 | 10 | 8 | 1.1 | 3 | 5 | n/a | n/a |
| Prostate adenoca. | 497 | 2.8 | 5 | 6 | 5 | 4 | 4 | 5 | 3 | | 0.262 |
| Skin cutaneous melanoma | 469 | 8 | 1.7 | 3 | 9 | 4 | 4 | 2.1 | 3 | 0.118 | 0.236 |
| Sarcoma | 259 | 5 | 10 | 6 | 8 | 5 | 8 | 4 | 5 | | |
| Testicular germ cell ca. | 150 | 2 | 7 | 7 | 9 | 5 | 6 | 3 | 3 | | 0.072 |
| Thymoma | 120 | 5 | 3 | 3 | 8** | 4 | 4 | 3 | 4 | 0.017 | 0.290 |
| Thyroid cancer | 501 | 3 | 2.4 | 4 | 6 | 3 | 3 | 3 | 5 | | |
| Uterine corpus endothelial carc. | 177 | 1.7 | 2.8 | 5 | 6 | 4 | 7 | 3 | 6 | | |
| **Overall %** | | **3.4** | **4.3** | **4.7** | **7.7** | **4.1** | **3.7** | **3.7** | **4.8** | | |

OS, Overall Survival, DFS, Disease/Progression Free Survival. Only p-values < 0.35 are shown. OS and DFS values describe negative impact on survival, unless marked by * (correlation with better survival). Expression alterations are predominantly observed as overexpression unless marked by ** (where downregulation is observed). SCC, squamous cell carcinoma.

Table S6. cBioPortal RNA-seq data (for cancers with provisional TCGA data): frequency of expression alterations (%) in genes involved in glutathione synthesis and correlation to survival (Logrank test p-value).

| Cancer type | n | GCLC | GCLM | GSS | GGCT | OPLAH | GSR | OS | DFS |
|---|---|---|---|---|---|---|---|---|---|
| Adrenocortical ca. | 79 | 5 | 1.3 | 6 | 16 | 6 | 4 | 0.093 | |
| Cholangiocarcinoma | 36 | 8 | 2.8 | 2.8 | 11 | 8 | 6 | | 0.296 |
| Bladder urothelial ca. | 408 | 4 | 4 | 18 | 16 | 11 | 2.9 | 6.327 e-3 | 0.231 |
| Colorectal adenoca. | 379 | 8 | 4 | 39 | 20 | 7 | 16** | | |
| Breast invasive ca. | 1093 | 5 | 7 | 10 | 8 | 15 | 4 | 0.017 | 0.339 |
| Glioblastoma mult. | 160 | 4 | 4 | 14 | 31 | 3 | 8 | 0.175 | 0.087 |
| Cervical SCC | 304 | 3 | 3 | 12 | 8 | 8 | 4 | | |
| Esophageal carcinoma | 184 | 7 | 4 | 13 | 16 | 11 | 8 | | |
| Stomach adenoca. | 415 | 7 | 6 | 14 | 13 | 15 | 9 | 0.04* | 0.163* |
| Uveal melanoma | 80 | 13 | 1.3 | 8 | 6 | 35 | 6 | | |
| HNSCC | 520 | 6 | 6 | 10 | 8 | 12 | 6 | 0.019 | 2.763e-3 |
| Kidney renal clear cell ca | 533 | 5 | 4 | 6 | 8 | 5 | 5 | 0.185 | 0.070 |
| Kidney renal papillary cell ca. | 290 | 4 | 4 | 16 | 7 | 8 | 3 | 0.051 | 0.041 |
| Liver hepatocellular ca. | 371 | 6 | 3 | 5 | 7 | 20 | 2.7 | | |
| Lung adenocarcinoma | 515 | 6 | 2.9 | 8 | 5 | 14 | 5 | 0.035 | |
| Lung SCC | 501 | 7 | 7 | 10 | 10 | 12 | 5 | | 0.211* |
| AML | 173 | 5 | 6 | 4 | 3 | 6 | 7 | | |
| Ovarian serous cystadenoca. | 307 | 4 | 4 | 13 | 4 | 35 | 3 | | 0.337* |
| Pancreatic adenoca. | 178 | 4 | 2.2 | 8 | 7 | 7 | 5 | 0.15* | |
| Mesothelioma | 87 | 7 | 3 | 6 | 3 | 9 | 6 | n/a | n/a |
| Prostate adenoca. | 497 | 6 | 4 | 4 | 8 | 11 | 6** | 0.13* | 7.141e-3 |
| Skin cutaneous melanoma | 469 | 13 | 5 | 11 | 18 | 8 | 8 | 2.585 e-3 | 0.184 |
| Sarcoma | 259 | 2.7 | 4 | 9 | 12 | 3 | 5 | 0.011 | 0.041 |
| Testicular germ cell ca. | 150 | 7 | 13 | 2.7 | 26 | 4 | 9 ** | 0.05 | |
| Thymoma | 120 | 6 | 3 | 7 | 6 | 4 | 5 | 1.111 e-4 | |
| Thyroid cancer | 501 | 2.4 | 1.8 | 6 | 4 | 1.4 | 1.6 | | |
| Uterine corpus endothelial carc. | 177 | 5 | 4 | 6 | 6 | 12 | 3 | | |
| **Overall %** | | **5.9** | **4.2** | **9.9** | **10.6** | **10.8** | **5.7** | | |

OS, Overall Survival, DFS, Disease/Progression Free Survival. Only p-values < 0.35 are shown. OS and DFS values describe negative impact on survival, unless marked by * (correlation with better survival). Expression alterations are predominantly observed as overexpression unless marked by ** (where downregulation is observed). SCC, squamous cell carcinoma.

Table S7. qRT-PCR primers' sequences.

| Gene | Forward (5'-3') | Reverse (5'-3') |
|---|---|---|
| *DNMT1* | GAGCCACAGATGCTGACAAA | GACACAGGTGACCGTGCTTA |
| *DNMT3A* | AAGGAGGAGCGCCAAGAG | GGATGGGGACTTGGAGATCA |
| *DNMT3B* | GGGAGGTGTCCAGTCTGCTA | GGCTTTCTGAACGAGTCCTG |
| *TP63* | GTTTCGACGTGTCCTTCCAG | TCTGGATGGGGCATGTCTTT |
| *KRT5* | TGAGGTCAAGGCCCAGTATG | ATCTCATGCTTGGTGTTGCG |
| *IVL* | AACACAAAGGGATCAGCAGC | GCTCCAACAGTTGCTCTTTCT |
| *PTGS2 (COX-2)* | TCATCATCAGCGCCCTCAA | GCTCGTTCACAGCCTTCATG |
| *PTGER1 (EP-1)* | GCCAGCTTGTCGGTATCATG | CTGCAGGGAGGTAGAGCTC |
| *PTGER2 (EP-2)* | AAGCTGTGGTCAAGGCTACA | GCCAAGTACCATGCTCACTG |
| *PTGER3 (EP-3)* | GGATCATGTGCGTGCTGTC | TGTGTCTTGCAGTGCTCAAC |
| *PTGER4 (EP-4)* | TGCTCATCTGCTCCATCCC | ATTCGGATGGCCTGCAAATC |
| *ALOX5* | ACATCTACCTCAGCCTCGTG | AGTTCCTCGTCCACAGTCAC |
| *ALOX15B* | AAATCAAGGGGTTGCTGGAC | AACTGGGAGGCGAAGAAGG |
| *ALOX12* | CTTGCTGAACACTCACCTGG | ATGGTGTAGCGGATATGGGG |
| *LTA4H* | GACTTCTGGGAAGGAACACC | TGCCACCAGTTCTTTAGGGA |
| *CYP2E1* | CGGAACTATGGGATGGGGAA | CGGAAGAGGATGTCGGCTAT |
| *ITGAM (CD11b)* | TGTTTCACGGAACCTCAGGA | ATCCATTGTGAGGTCCTGGC |
| *ACTB* | AAAGACCTGTACGCCAACAC | GTCATACTCCTGCTTGCTGAT |

A

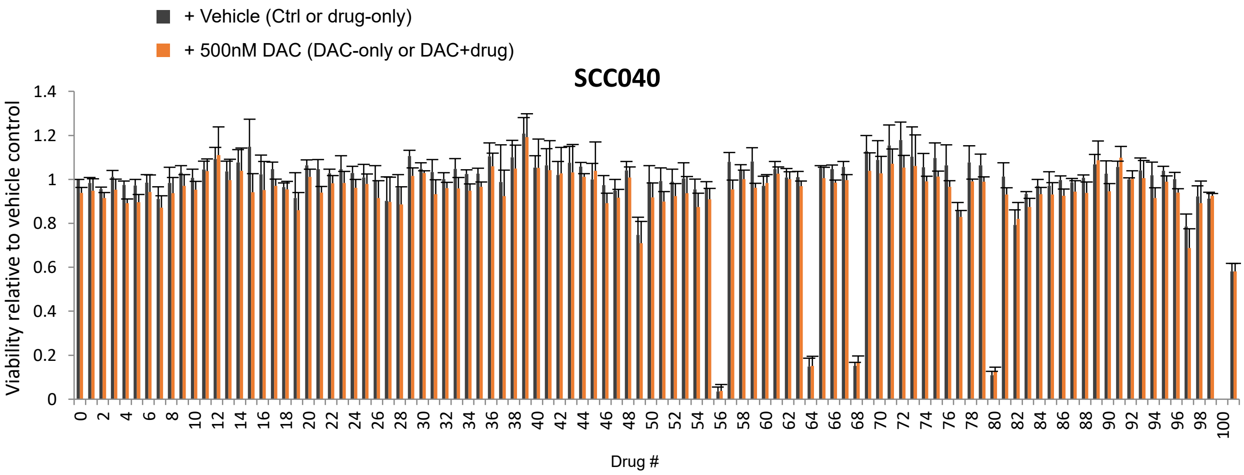


B

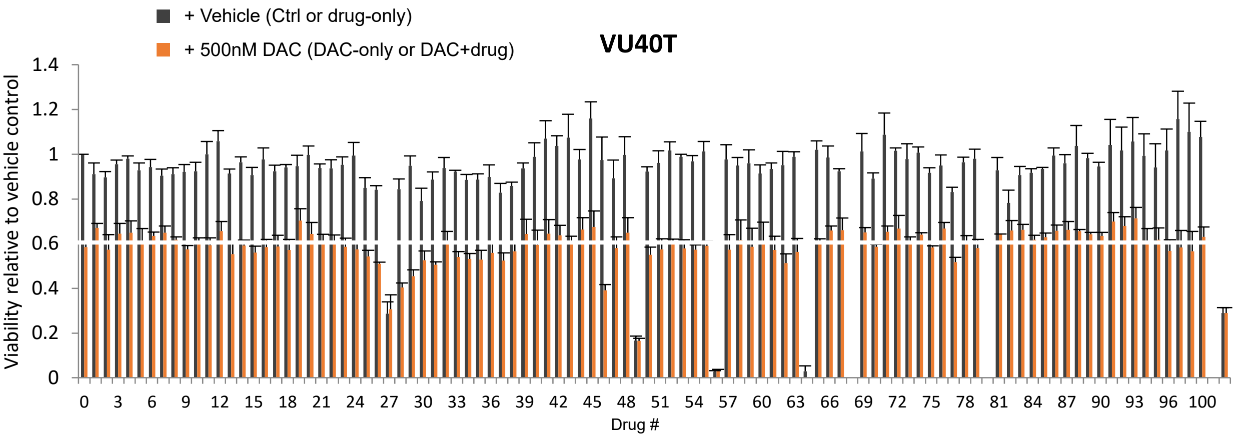


**Figure S1. Sensitivity of HNSCC cells to DAC treatment can be increased by drug combinations.**

**A-B.** DAC sensitizing assay: SCC040 (**A**) and VU40T (**B**) cells were subjected to 96h treatment with one of a panel of 100 drugs, +/-500 nM DAC. Viability was recorded and is shown here relative to the vehicle only control cells (Drug #0 black bar). The horizontal white line shows the effect of 500 nM DAC alone. The bars far right (Drug #101) show the effect of 10 µM DAC. The sensitizing effect is observed when the combined effect of the two drugs is more effective than both DAC alone and the drug alone. The assay was performed in triplicate and error bars represent SEM.
Drug #28: zinc acetate, #29: valproic acid, #46: paracetamol.

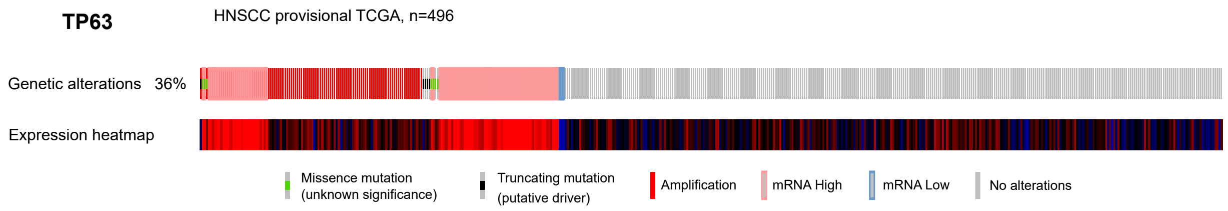


**Figure S2 - *TP63* gene alterations in head and neck squamous cell carcinoma.**
Genomic and expression alterations in *TP63* gene in 496 tumours from the provisional TCGA cohort of HNSCC patients. Upper graph shows genetic and expression (z-score => or =<2.0) alterations, explained in the legend; lower graph depicts expression heatmap.

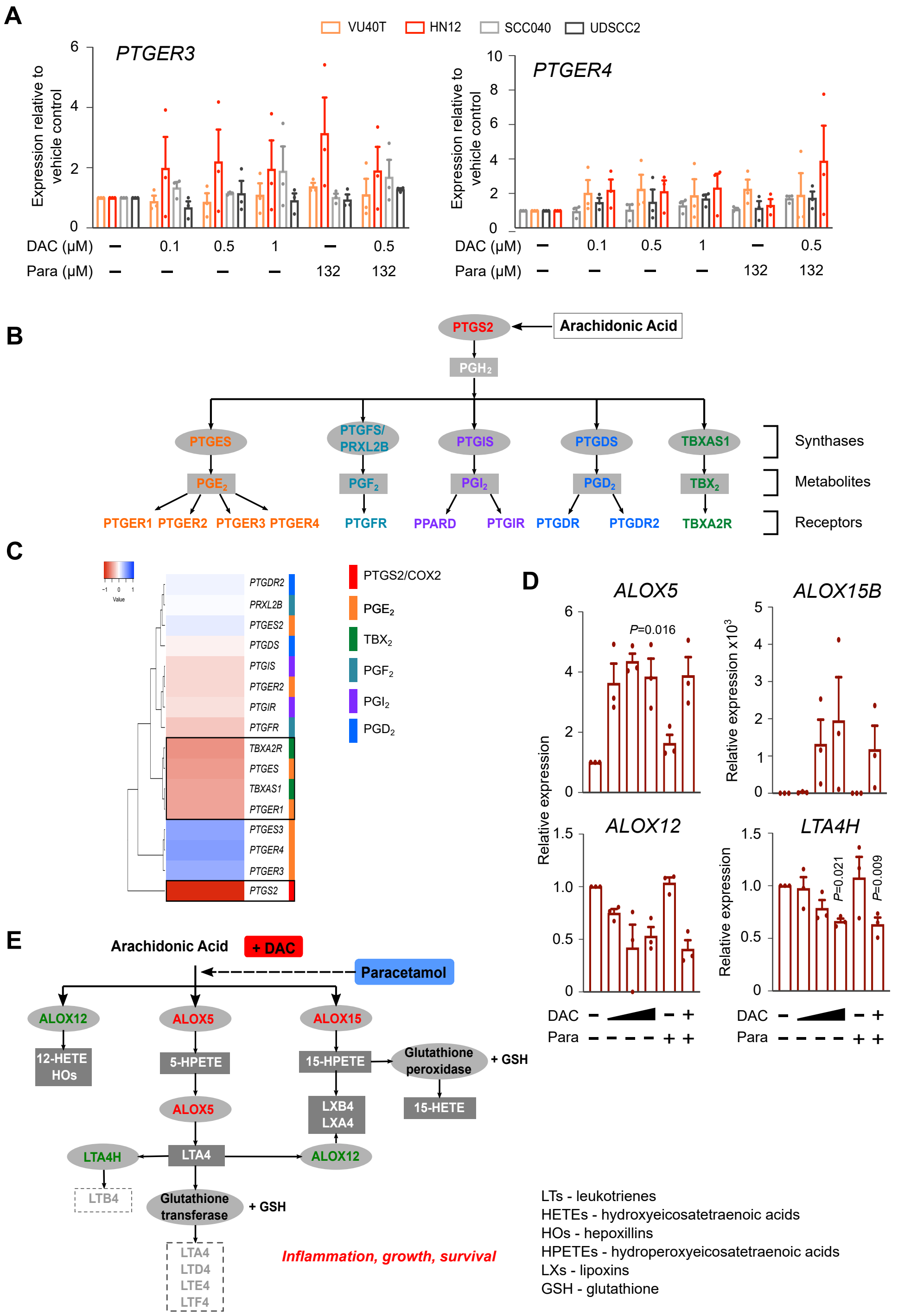


**Figure S3 - COX-2 pathways are selectively affected by DAC treatment.**

**A.** qRT-PCR for *PTGER3* and *PTGER4* genes in HNSCC cell lines treated for 96h with DAC and/or paracetamol as indicated. Data are shown as relative to vehicle control.

**B.** Schematic representation of the cyclooxygenase pathway. COX enzymes convert arachidonic acid (AA) into prostaglanding $H_2$ which is then converted to prostanoids (prostaglandins $PGE_2$, $PGF_2$, $PGI_2$, $PGD_2$ and thromboxane $TXA_2$) by respective synthases. Prostanoids are then recognized by G-protein coupled receptors in both autocrine and paracrine manner.

**C.** DAC-induced expression fold changes for genes encoding synthases and receptors involved in all cyclooxygenase pathways. Data obtained through RNA-seq in VU40T cells treated with 500 nM DAC for 96h. Black boxes indicate genes with highest upregulation. Colour-coded association with a specific pathway is shown to the right.

**D.** qRT-PCR for LOX pathway enzymes: *ALOX5*, *ALOX15B*, *ALOX12* and *LTA4H* in VU40T cells treated for 96h as indicated. Data are shown as relative to vehicle control.

**E.** Schematic of LOX pathway with confirmed DAC effects (up-regulation in red, down-regulation in green). Leukotrienes (LTs) shown in dotted boxes were below the detection levels of ELISA.

Data information: In A and D n=3, for each cell line a matched One-Way ANOVA with Dunnett's correction to compare all treatments to Ctrl. Values displayed as means +/-SEM. Only significant p-values are shown.

**A**

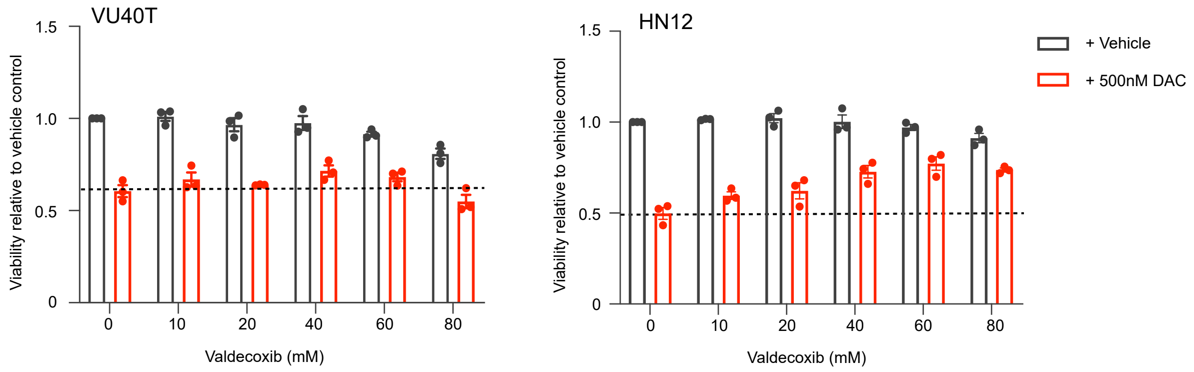


**B**

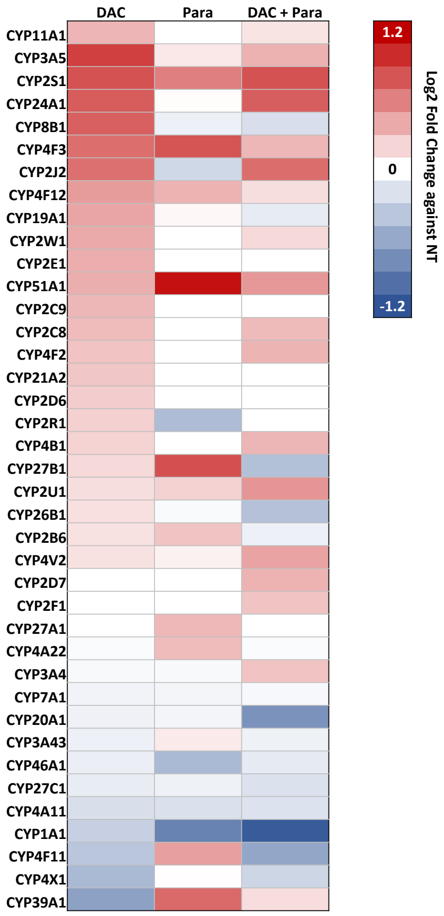


**Figure S4. The mechanisms of DAC-paracetamol synergy are specific to paracetamol and involve CYP enzymes upregulation**

**A.** Valdecoxib does not sensitize HNSCC cells to DAC treatment. VU40T and HN12 cell viability after 96h of treatment with indicated concentrations of Valdecoxib with or without 500 nM DAC. Dotted lines indicate viabillity at 500 nM DAC only. Mean ±SEM; n=3.

**B.** Gene expression changes (RNA-seq data) in VU40T cells of CYP enzymes shown as heatmap of $\log_2$ fold change values after indicated treatments (DAC, paracetamol, DAC+paracetamol) against untreated control.

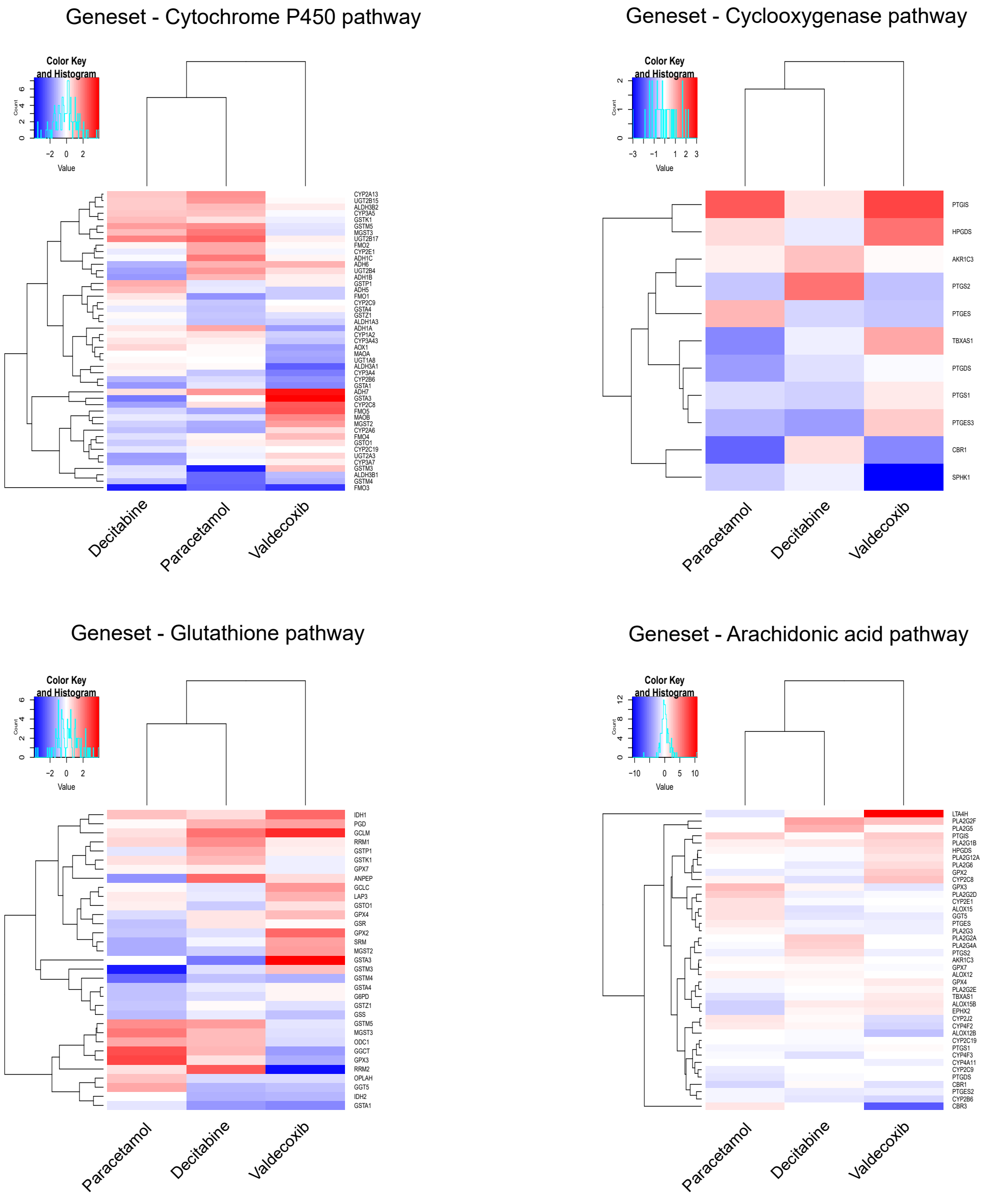


**Figure S5 - DAC and paracetamol gene expression signatures share similar pathway enrichment**. Drug perturbation signatures of Decitabine, paracetamol and valdecoxib, plotted for subsets of genes representing key pathways of interest that were identified from the DSEA analysis. Genes pertaining to each pathway were subsequently obtained from the corresponding genesets identified in MSigDB collection. Clustering of the drug perturbation profiles across these pathways indicates that Decitabine and paracetamol share similar drug perturbation profiles, compared to valdecoxib.

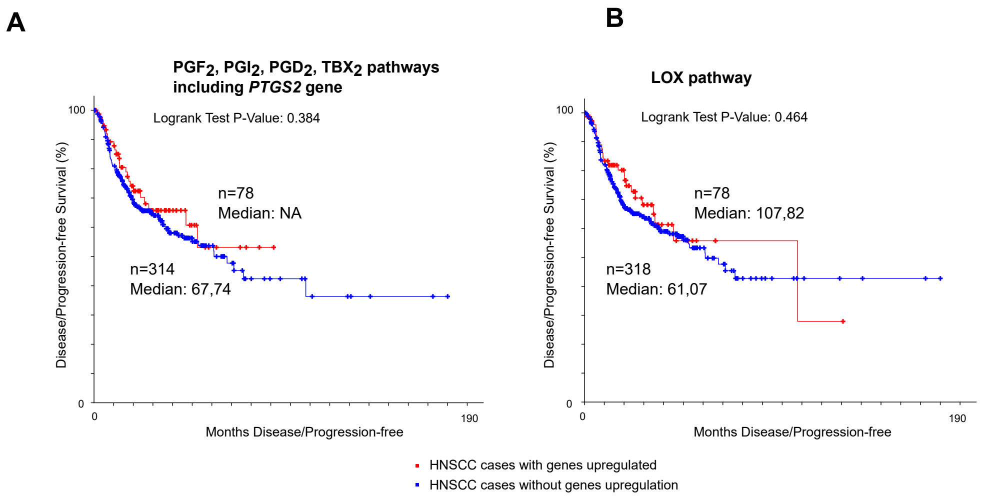


**Figure S6 - COX-2 pathway alterations selectively affect HNSCC patients survival.**

**A.** Disease/Progression-free survival curves for 392 HNSCC patients with data available in TCGA provisional cohort and gene upregulation for cyclooxygenase pathways other then $PGE_2$ (but including *PTGS2* gene).

**B.** Disease/Progression-free survival curves for 392 HNSCC patients with data available in TCGA provisional cohort and gene upregulation for LOX pathway.

Kaplan-Meier Estimates were obtained and plotted using cBioPortal.

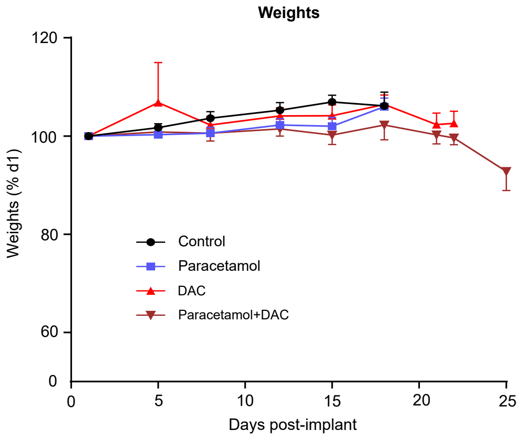


**Figure S7 - *In vivo* potential of DAC-paracetamol combination.**
Mice weights recorded during the efficacy study comparing treatments with DAC alone, paracetamol alone and DAC+paracetamol combination.
